# Transient Partial Inhibition of DNA Polymerase α Promotes Chromatin Plasticity and Enhances Human Cell Reprogramming

**DOI:** 10.64898/2026.10.25.761701

**Authors:** Jian Qi Luo, Yanfeng Yang, Guanghui Yang, Ryan Lee, Binbin Ma, Xin Chen

**Author notes:** Corresponding Authors: Binbin Ma, Ph.D., Xin Chen, Ph.D. Equal contributions (Jian Qi Luo, Yanfeng Yang, Guanghui Yang).

## Abstract

Somatic cell reprogramming requires extensive remodeling of epigenomic and transcriptional state, yet the mechanisms that enhance chromatin plasticity during cell-fate conversion remain incompletely understood. Beyond its essential role in genome duplication, DNA replication provides a dynamic window for chromatin remodeling and may therefore influence cellular plasticity during reprogramming. Here, we show that low-dose, transient inhibition of the lagging-strand DNA polymerase α (Polα) complex markedly enhances reprogramming of human peripheral blood mononuclear cells into induced pluripotent stem cells (iPSCs). Fine-tuned inhibition of the catalytic subunit PolA1 increases cell survival under reprogramming-associated stress and enhances cellular resilience to diverse external stresses, preserves genomic stability, and accelerates the acquisition of pluripotent features. PolA1-inhibited cells undergo rapid increases in chromatin accessibility following reprogramming factor delivery and display accelerated progression toward the iPSC state. Consistent with enhanced developmental plasticity, colonies derived from PolA1 inhibitor-treatment display dome-shaped morphology, increased expression of several naïve pluripotency markers such as SUSD2. Integrated imaging and single-nucleus multiomic analyses establish transient PolA1 inhibition as a potent regulator of early chromatin remodeling during reprogramming. Functionally, PolA1 inhibitor-derived iPSC lines maintain stable pluripotency and exhibit enhanced differentiation potential, including increased embryoid body budding and accelerated cardiomyocyte differentiation and maturation. Notably, transient and low-dose PolA1 inhibition alters replication-coupled histone incorporation without reducing DNA replication speed or affecting cell cycle progression. This two-step regime primes human PBMCs for sustained gains in reprogramming efficiency with pre- and post-treatment. Together, these findings identify replication-coupled chromatin regulation as an underappreciated mechanism regulating human cell fate plasticity and demonstrate that transient modulation of Polα can simultaneously enhance epigenomic plasticity and preserve genome stability during reprogramming.

## Introduction

Cellular reprogramming of differentiated somatic cells into induced pluripotent stem cells (iPSCs) has transformed regenerative medicine by enabling patient-specific disease modeling, drug discovery, and cell-based therapies^1–3^. Despite its broad potential, reprogramming remains inefficient because differentiated cells must overcome multiple barriers to erase somatic identity and establish a pluripotent state^4,5^. Numerous strategies have therefore been developed to improve reprogramming efficiency by manipulating transcription factors, signaling pathways, microRNAs, metabolism, and chromatin-modifying enzymes^6–10^. However, the mechanisms that enable cells to overcome epigenetic barriers and acquire a highly plastic chromatin state remain incompletely understood.

Successful reprogramming requires extensive remodeling of chromatin and epigenetic information. DNA replication provides a critical opportunity for this process because newly synthesized DNA must be assembled into chromatin through the recycling of parental histones, incorporation of newly synthesized histones, and re-establishment of epigenetic information.

Therefore, DNA replication is tightly coupled to chromatin organization, epigenetic inheritance, and transcriptional regulation. These connections raise the possibility that the replication machinery may influence cellular plasticity beyond its canonical role in genome duplication. However, whether controlled modulation of DNA replication can facilitate cell-fate conversion remains largely unexplored.

DNA replication is inherently asymmetric, with leading- and lagging-strand synthesis relying on distinct enzymatic and regulatory mechanisms that may differentially influence genome maintenance and epigenetic inheritance. The eukaryotic RNA Primase–DNA polymerase α (Polα) complex is essential for initiating lagging-strand synthesis. Primase generates short RNA primers that are extended by the catalytic PolA1 subunit to initiate synthesis of Okazaki fragments. Because Primase–Polα activity is required repeatedly on the lagging strand, this complex is positioned at the interface between DNA synthesis and replication-coupled chromatin assembly. Notably, Polα has been reported to facilitate recycling of parental histones to the lagging strand during replication^11^. Because parental histone recycling contributes to the preservation of epigenetic information, we hypothesized that modest, transient modulation of Polα activity could alter replication-coupled chromatin assembly and thereby facilitate chromatin remodeling required for cell-fate transitions while maintaining sufficient replication capacity.

Our previous studies in invertebrates support this hypothesis: We discovered that developmental regulation of lagging-strand DNA replication machinery programs cell-type-specific epigenomes in the *Drosophila* male germline through replication-coupled histone incorporation^12^. Differentiating progenitors express approximately twofold higher levels of Polα than germline stem cells, whereas reducing *primase1* dosage or PolA1 activity using an inhibitor induces a stem cell-like pattern of histone incorporation in progenitor cells. These findings indicate that lagging-strand replication machinery can modulate cell-type-specific chromatin states in an adult stem cell lineage. Consistently, our subsequent studies found that reducing the dosage of Polα complex components enhances stem cell maintenance, regeneration, and cellular reprogramming in multiple stem cell lineages in *Drosophila* and *C. elegans*^13^. Together, these findings in invertebrates suggest that controlled modulation of replication-coupled chromatin assembly can influence cell-state plasticity and tissue resilience.

Here, we investigate whether this principle extends to human somatic cell reprogramming and explore the mechanisms underlying the effect. Using peripheral blood mononuclear cells (PBMCs) as a clinically relevant starting population, we show that fine-tuned, transient inhibition of PolA1 during the early stages of reprogramming markedly enhances establishment of stable iPSC lines while increasing resilience to reprogramming-associated and external stress. Integrated imaging and single-nucleus multiomic analyses reveal that transient PolA1 inhibition rapidly increases chromatin accessibility and accelerates acquisition of pluripotent identity.

PolA1 inhibition-derived iPSC lines maintain stable pluripotency and exhibit enhanced differentiation potential, supporting a more developmentally plastic state. These findings identify replication-coupled chromatin regulation as an underappreciated mechanism controlling human cell fate plasticity and establish transient modulation of Polα activity as a promising strategy of enhancing the efficiency and genome stability of human iPSC generation.

## Results

### Transient, low-dose PolA1 inhibition enhances the efficiency and kinetics of human PBMC reprogramming into iPSCs

To investigate the effects of Polα inhibition in human cells, we used a pharmacological inhibitor that blocks the DNA-binding and primer elongation activities of the catalytic PolA1 subunit (PolA1i)^14,15^.We first optimized PolA1i dosing in human PBMCs, which provide a clinically relevant starting population for reprogramming because they can be readily obtained through minimally invasive blood collection. Higher-dose PolA1 inhibition (1 μM) markedly impaired DNA synthesis, as indicated by reduced EdU incorporation (Fig. S1A). In contrast, lower doses (10 and 100 nM) largely preserved DNA synthesis capacity, with EdU incorporation comparable to vehicle-treated controls (Fig. S1A). Notably, 100 nM PolA1i modestly increased the S-phase index (Fig. S1B), likely due to a prolonged S phase rather than a substantial reduction in DNA synthesis^14,16^. Based on these findings, we selected 10 and 100 nM PolA1i for initial reprogramming experiments to determine whether low-dose Polα inhibition could enhance cellular plasticity while preserving DNA replication capacity.

PBMCs were reprogrammed by Sendai virus-mediated expression of OCT3/4, SOX2, KLF4, and c-MYC^17,18^. Based on our previous finding that a brief exposure to an optimized concentration of PolA1i (10uM; 4 hours) was sufficient to alter replication-coupled histone incorporation patterns in the *Drosophila* male germline^12^, we hypothesized that transient PolA1 inhibition during the early stages of PBMC reprogramming could influence cell fate conversion. To test this possibility, PBMCs were treated with PolA1i for two days before viral transduction and for an additional three days following transduction (Fig. 1A). Compared with vehicle-treated controls, PolA1i-treated PBMCs produced substantially more colonies positive with SSEA-4 (Stage-Specific Embryonic Antigen-4), an early cell-surface marker associated with pluripotency^17^ (Fig. 1B), with increased colony numbers and colony sizes detectable as early as day 9 and sustained through day 16 (Fig. 1C-E). Across six independent donor samples, 10 nM PolA1i increased reprogramming efficiency by approximately threefold on average, demonstrating a robust and reproducible effect across donor samples. The 100 nM dose also increased colony formation but exhibited greater donor-to-donor variability (Fig. 1C), potentially reflecting its stronger effects on S-phase progression (Fig. S1).

**Figure 1:**
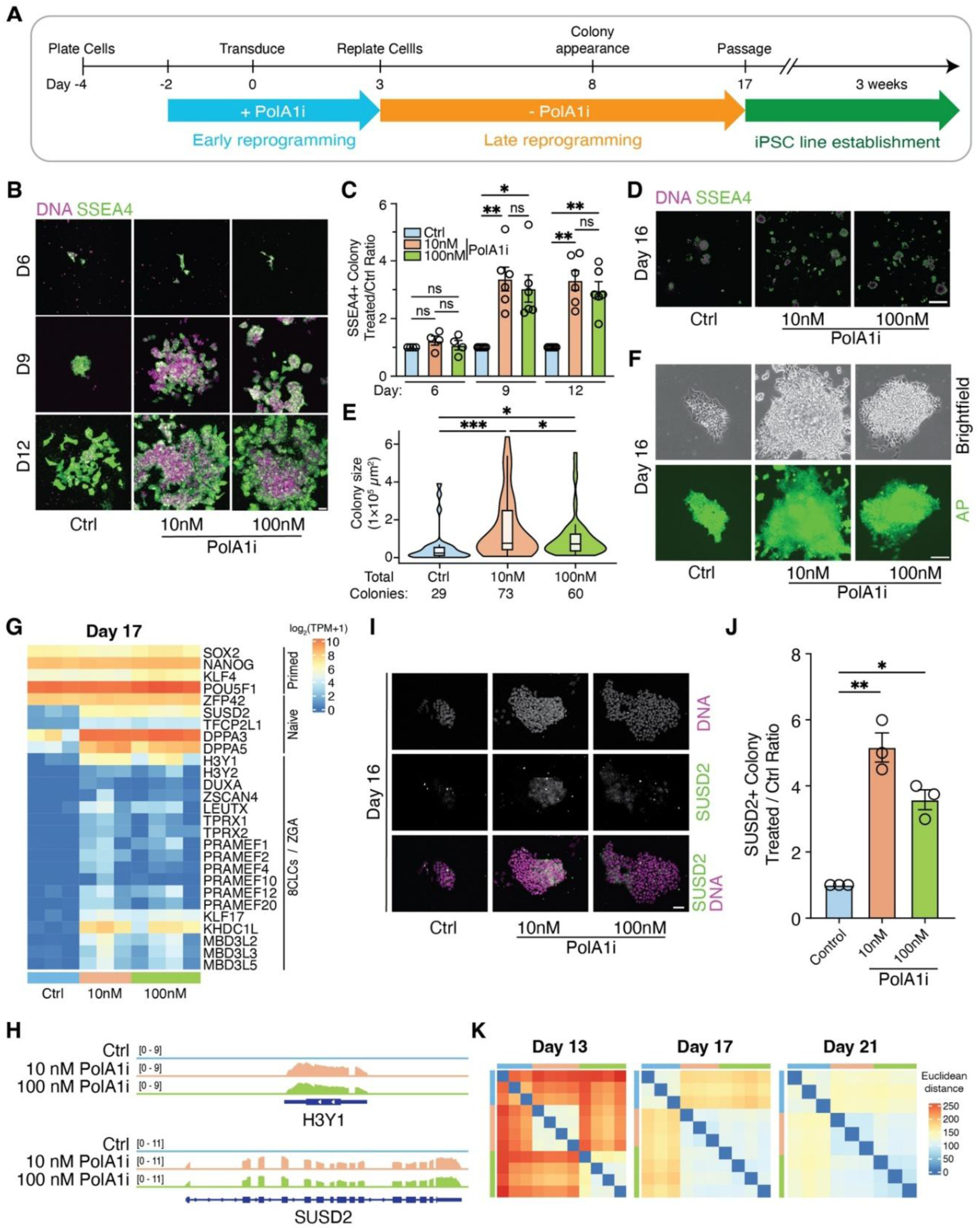
Transient, low-dose PolA1 inhibition enhances the efficiency and kinetics of human PBMC reprogramming into iPSCs. (**A**) Schematic overview of the PBMC-to-iPSC reprogramming protocol with or without transient PolA1 inhibition. Representative immunofluorescence images (**B**) of SSEA-4-positive colonies and quantification (**C**) of SSEA4-positive cells from day 6 to day 12 in control and PolA1 inhibitor-treated conditions. Representative tiled image of SSEA4-positive colonies (**D**) and quantification (**E**) on day 16. (**F**) Representative alkaline phosphatase staining of control and PolA1 inhibitor-treated colonies. (**G**) Heatmap showing changes in the expression of pluripotency-state-associated marker genes in reprogrammed iPSCs on day 17 before passaging. (**H**) Representative RNA-seq genome browser tracks for SUSD2 and H3Y1. Representative immunofluorescence images of SUSD2-positive colonies (**I**) and quantification (**J**) on day 16. (**K**) Hierarchical clustering of RNA-seq data from control and PolA1 inhibitor-treated colonies collected on days 13, 17, and 21. Scale bars: 50 µm (**B**), 2000 µm (**D**), 100 µm (**F**), 50 µm (**I**).

PolA1i-treated colonies retained key features of pluripotency. Alkaline phosphatase staining confirmed the formation of pluripotent colonies (Fig. 1F), and single-colony transcriptomic profiling revealed comparable expression of core pluripotency markers, including SOX2 and NANOG, between control and PolA1i-derived iPSCs (Fig. 1G). Notably, PolA1i-treated colonies exhibited a more compact, dome-like morphology and increased expression of naïve pluripotency-associated genes, including SUSD2, TFCP2L1, DPPA3, and DPPA5^19–21^ (Fig. 1F, 1I-J). These colonies also showed elevated expression of genes associated with the 8-cell-like cell (8CLC) state and zygotic genome activation (ZGA), including H3Y1, ZSCAN4, and LEUTX especially in colonies on day 17^22^. Together, these findings demonstrate that transient PolA1 inhibition enhances PBMC reprogramming efficiency while promoting the acquisition of molecular features associated with a more naïve and developmentally plastic pluripotent state.

To determine whether PolA1 inhibition alters the dynamics of reprogramming, we next compared transcriptomic heterogeneity across multiple time points (Fig. 1K). Hierarchical clustering revealed substantial variability among day 13 control colonies, consistent with asynchronous progression through the early stages of reprogramming. In contrast, colonies treated with 10 nM PolA1i exhibited more coherent transcriptional profiles and clustered more closely together, indicating greater synchronization of their reprogramming trajectories. The 100 nM condition also reduced transcriptional heterogeneity relative to controls, although this effect was less consistent. By day 17, control colonies became progressively more homogeneous but remained transcriptionally distinct from PolA1i-treated colonies. By day 21, however, all groups largely converged, with minimal residual transcriptional differences. These findings indicate that transient PolA1 inhibition reduces transcriptional heterogeneity during the early phase of reprogramming and promotes more coordinated establishment of pluripotency-associated gene expression programs.

Together, these findings demonstrate that transient low-dose PolA1 inhibition reshapes the early trajectory of PBMC reprogramming by enhancing colony formation, reducing transcriptional heterogeneity, and facilitating the emergence of naïve pluripotency- and 8CLC-associated molecular states. Because the 10 nM PolA1i condition produced the most robust and reproducible effects, we used this dose for subsequent studies.

### PolA1i-derived iPSC lines maintain stable pluripotency and exhibit enhanced differentiation capacity

Individual iPSC colonies were manually isolated and expanded to establish independent cell lines. By passage 2, PolA1i-derived lines (Treated) exhibited a compact, dome-shaped colony morphology, whereas control-derived lines (Ctrl) were generally flatter (Fig. 2A).

**Figure 2.**
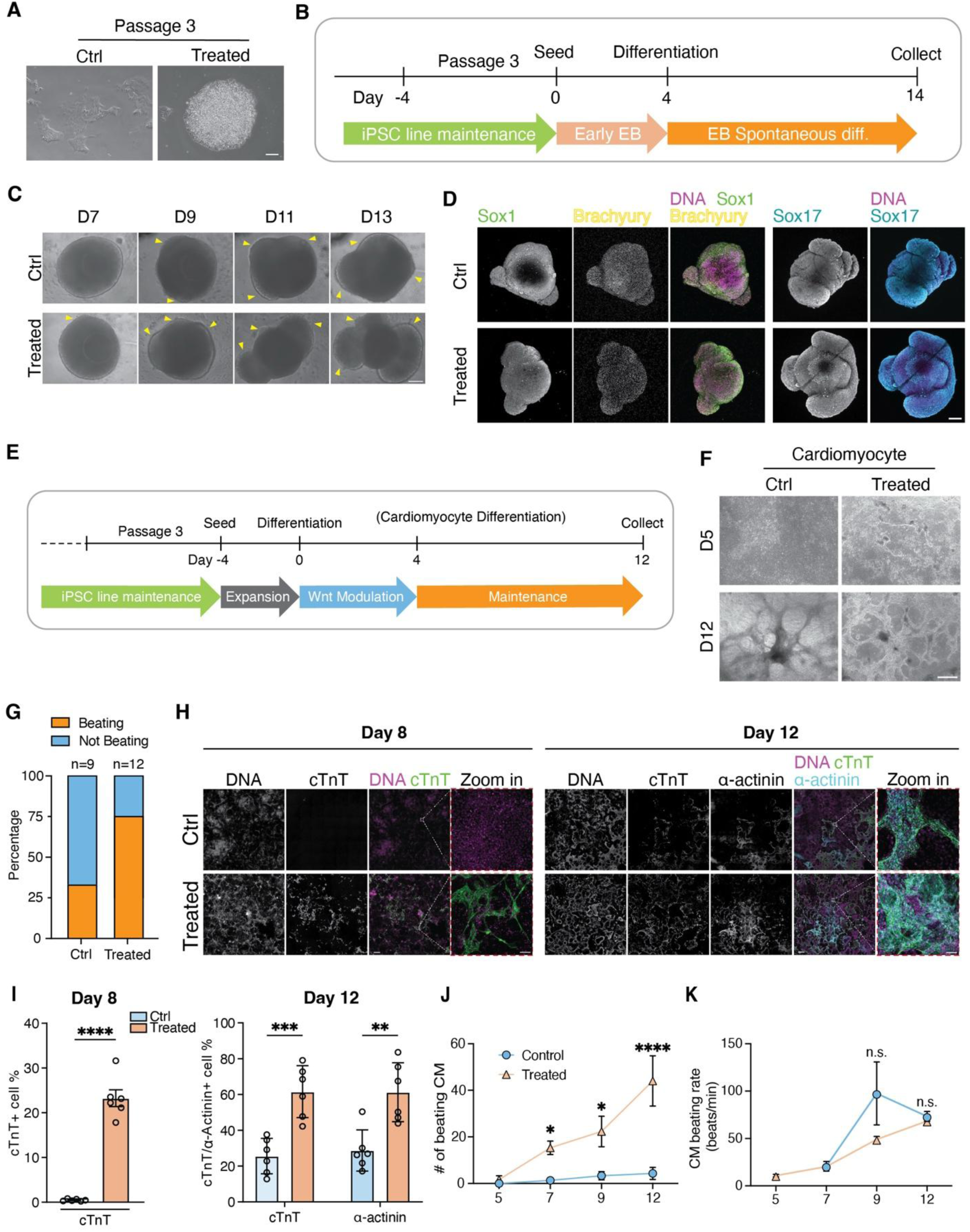
Early passage of PolA1i-derived iPSC lines maintain stable pluripotency and exhibit enhanced differentiation capacity. (**A**) Representative bright-field images of passage-3 iPSC lines derived from control- or PolA1i-treated cells. (**B**) Schematic overview of the re-differentiation protocol of iPSC generated with or without transient PolA1 inhibition toward embryoid body formation. (**C**) Representative bright-field images of passage-3 iPSC-derived EBs with buds during differentiation from day 7 to day 13. (**D**) Representative immunofluorescence images showing expression of the ectodermal marker SOX1, mesodermal marker brachyury, and endodermal marker SOX17 in EBs generated from control- and PolA1i-derived iPSC lines. (**E**) Schematic overview of the directed differentiation protocol of iPSC generated with or without transient PolA1i inhibition into cardiomyocyte lineage. (**F**) Representative bright-field images of cardiomyocyte differentiation on days 5 and 12 and quantification of the number of beating colonies in control- and PolA1i-derived cultures(**G**). Representative immunofluorescence images of cardiac troponin T (cTnT) and α-actinin in day 12 cardiomyocytes (**H)** and corresponding quantification (**I)**. Quantification of the number of emerging beating colonies (**J**) and beating rate (**K**) during differentiation. Scale bars: 100 μm (**A**), 200 μm (**C**), 200 μm (**D**), 500 μm (**F**), 1000 μm (**H**) and 50 μm (**H-Zoom in**).

Although some Ctrl lines became more compact with passaging, only 2 of 6 Ctrl lines displayed dome-shaped colonies at passage 10, compared with 6 of 10 Treated lines (Fig. S2A). Both cell lines exhibited the maintenance of the pluripotent stem cell markers such as Nanog, Oct-4, and Sox2 (Fig. S2B). Overall, iPSC lines passaged from treated iPSCs more frequently exhibited compact, dome-shaped colonies that resemble of mouse embryonic stem cells, prompting us to further investigate whether the PolA1i treatment enhances developmental plasticity^23,24^, with a focus on earlier passaging iPSC lines.

We next assessed the differentiation potential of these iPSC lines using spontaneous embryoid body (EB) formation (Fig. 2B-D) and directed cardiomyocyte differentiation^25–28^(Fig. 2E-J). During spontaneous EB differentiation (Fig. 2B), passage-3 of Treated lines began to form buds by day 11, with bud formation increasing in both Ctrl and Treated lines by day 13 (Fig. 2C). A consistent trend of higher number of EB buds formation in Treated lines than in Ctrl lines throughout differentiation suggests a potential enhanced differentiation and cellular plasticity in PolA1i-derived iPSCs during differentiation progress^29^ (Fig. 2E, S2C). To assess multilineage differentiation and pluripotency of early passage iPSC lines, we immunostained EBs for markers of the three germ layers (Fig. 2D). EBs generated from both Ctrl and Treated lines expressed markers of endoderm, mesoderm, and ectoderm, confirming the pluripotency and multilineage differentiation potential of both iPSC lines.

We next assessed directed differentiation using an established WNT-modulating small-molecule protocol for cardiomyocyte generation^30^ (Fig. 2E). By day 5, Treated cultures exhibited organized cardiomyocyte-like fibers, whereas Ctrl culture did not (Fig. 2F). Immunostaining for cardiac troponin T (cTnT) and a-actinin confirmed cardiomyocyte differentiation in both Ctrl and Treated iPSCs^30^ (Fig. 2H). At day 8, Treated cultures contained a higher proportion of cTnT-positive cells comparing to Ctrl, whereas by day 12, a significantly higher percentage of cells in Treated groups expressed both cTnT and a-actinin markers than Ctrl group (Fig. 2I). To determine whether early-passage iPSC lines could support efficient cardiomyocyte differentiation, we assessed spontaneous beating following differentiation. At passage-3, spontaneous beating was observed in 9 of 12 Treated lines compared with 3 of 9 Ctrl lines (Fig. 2G). Consistent with the earlier acquisition of cardiomyocyte features, spontaneous beating was detected in some Treated cultures by day 5, whereas no beating was observed in Ctrl cultures at this time point (Fig. 2J). By day 12, Ctrl cultures also exhibited spontaneous beating, however, the total beating colony count was approximately eightfold greater in Treated cultures than in Ctrl cultures (Fig. 2J). Finally, contraction frequency in Treated cardiomyocytes was comparable to that of cardiomyocytes generated from established later-passage iPSC lines, indicating that early-passage PolA1i-derived iPSC can undergo functional cardiomyocyte differentiation (Fig. 2K).

Together, these findings indicate that transient PolA1 inhibition during reprogramming promotes the establishment of iPSC lines with stable pluripotent features and enhanced differentiation potential. PolA1i-derived iPSCs exhibited promising EB differentiation, accelerated acquisition of cardiomyocyte features, and robust functional differentiation even at early passage comparing to Ctrl lines, suggesting that transient low-dose PolA1 inhibition during reprogramming establishes iPSC lines with enhanced developmental competence.

### Transient, low-dose PolA1 inhibition promotes reprogramming and alters chromatin accessibility and transcriptional states

Replication-coupled histone incorporation is closely linked to the restoration of chromatin structure after DNA replication and may therefore influence chromatin accessibility and gene expression. Having found that PolA1 inhibition selectively alters replication-coupled histone incorporation, we asked whether this change is accompanied by broader alterations in chromatin and transcriptional states during reprogramming.

PBMCs contain diverse cell populations, and reprogramming proceeds asynchronously through heterogeneous intermediate states. Bulk genomic or transcriptomic analyses could therefore average across distinct populations and obscure state-specific effects of PolA1 inhibition. We therefore performed single-nucleus (sn) multiomic profiling, which simultaneously measures gene expression (snRNA-seq) and chromatin accessibility (snATAC-seq) in the same nuclei. This approach allowed us to resolve both the starting cell populations and the reprogramming states, and to ask how PolA1 inhibition influences chromatin and transcriptional states along distinct reprogramming trajectories.

To investigate how PolA1 inhibition (PolA1i) promotes reprogramming, we collected control and PolA1i-treated samples at days 0, 3, and 7, spanning the early stages of reprogramming (Fig. 1A). We removed ambient RNA, doublets, and low-quality nuclei with insufficient RNA or ATAC reads. The filtered RNA and ATAC datasets from each sample were of high quality (Fig. S3A-B) and were integrated for joint analysis. UMAP embedding of the integrated multiomic data resolved 42 clusters (Fig. S3C), which we assigned to 13 cell types based on marker gene expression (Fig. 3A, S3D). Pseudotime analysis with Monocle3 identified a trajectory along which a small fraction of cells progressed toward an iPSC-like state, consistent with the cell type annotations (Fig. 3B). Consistent with these annotations, most cells retained PBMC identity, as shown by expression of the pan-PBMC marker PTPRC (Fig. 3C) and other lineage markers (Fig. S3D). A subset of cells, however, had begun expressing pluripotency factors, including POU5F1 (OCT4), SOX2, and KLF4 (Fig. 3C, S3D-E). These reprogramming intermediates lacked detectable expression of DPPA2 and ZFP42, indicating an incompletely reprogrammed state (Fig. S3D). Nearly all cells in this population originated from day 7 samples.

**Figure 3.**
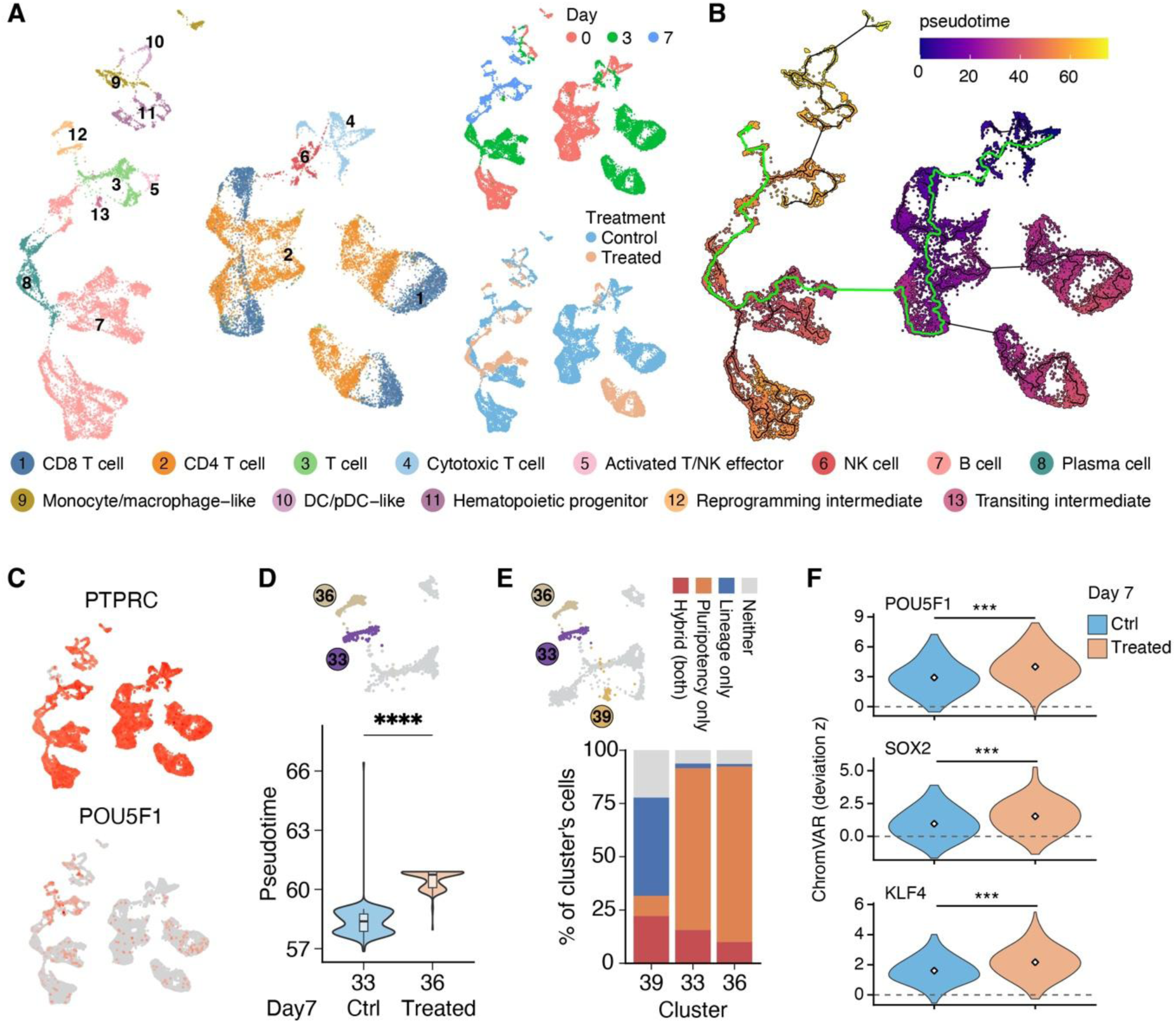
Single-nucleus multiomic profiling shows that PolA1 inhibition accelerates reprogramming. (**A**) Weighted-nearest-neighbor (WNN) UMAP of integrated snRNA-seq and snATAC-seq data from PBMCs undergoing reprogramming, colored by annotated cell type (left), sampling day (day 0, 3, or 7; top right), and treatment (Control or PolA1i; bottom right). Cell type annotations (1–13) are listed below the panel. (**B**) Monocle3 pseudotime trajectory on the same embedding, colored by pseudotime. Black lines show the principal graph, and the green line highlights the path toward the reprogramming intermediates. (**C**) Feature plots of PTPRC (pan-PBMC marker) and POU5F1 (OCT4) expression. Color intensity indicates normalized expression; gray cells have no detectable expression. (**D**) Pseudotime distribution of day 7 cells in cluster 33 (Control) and cluster 36 (PolA1i). Insets show the location of these clusters on the UMAP. Violin plots show the full distribution, and box plots show the median and interquartile range. ****P < 0.0001, two-sided Wilcoxon rank-sum test. (**E**) Composition of clusters 39 (transitioning intermediates), 33, and 36 (reprogramming intermediates) according to expression of pluripotency and lineage marker genes. Cells are classified as “pluripotency only,” “lineage only,” “hybrid” (both), or “neither”. Insets show the location of these clusters on the UMAP. (**F**) ChromVAR deviation z-scores for POU5F1, SOX2, and KLF4 motifs in day 7 control and PolA1i-treated cells. Diamonds mark the mean, and the dashed line Scores were centered on the median score of all motifs in each cell, so 0 (dashed line) indicates accessibility of a typical motif. ***P < 0.001, two-sided Wilcoxon rank-sum test.

Mapping the reprogramming intermediates back onto the original 42 clusters identified two constituent clusters, 33 and 36, which consisted predominantly of day 7 control and PolA1i-treated cells, respectively. Although both clusters represent reprogramming intermediates, PolA1i-treated cells in cluster 36 occupied a significantly more advanced position along the pseudotime trajectory than control cells in cluster 33 (Fig. 3D). This result suggests that PolA1i accelerates progression toward the iPSC-like state, consistent with the earlier emergence of iPSC colonies in treated cultures (Fig. 1B-C). Cluster 39, which lies adjacent to the reprogramming intermediates, comprised “transitioning intermediates.” In this population, only a minority of cells expressed pluripotency factors, whereas most retained PBMC lineage markers (Fig. S3D). Comparing cluster 39 with clusters 33 and 36 supports a model in which PBMC reprogramming proceeds through gradual silencing of lineage-specific genes accompanied by activation of pluripotency-associated genes (Fig. 3E). Notably, cluster 36 also showed increased chromatin accessibility at genomic regions containing POU5F1, SOX2, and KLF4 binding motifs, and these increases were more pronounced than the corresponding increases in gene expression (Fig. 3F, S3E). This pattern suggests that chromatin remodeling may contribute more than transcriptional activation to the PolA1i-driven advance, consistent with the permissive chromatin state of iPSCs after reprogramming.

A closer examination of cluster 39 showed that the cells expressing pluripotency genes came mainly from day 3 samples, where PolA1i-treated cells again showed an advantage (1% of cells) over control cells (0.5% of cells) (Fig. S3F). Together, these results support a continuous reprogramming trajectory in which pluripotency features emerge in a very small population of cells as early as day 3 after induction. PolA1 inhibition accelerates this process, and the acceleration is associated with higher expression of pluripotency genes and greater chromatin accessibility at their binding motifs.

### Transient, low-dose PolA1 inhibition promotes cell survival and suppresses cell-death programs during reprogramming

snRNA-seq analysis of day 3 samples revealed that low-dose PolA1 inhibition increased the expression of anti-apoptotic genes. Notably, a larger fraction of PolA1i-treated cells had detectable transcripts of *MCL1, BCL2L1, BCL2A1, BIRC5*, and *XIAP*, with a smaller increase in *BCL2* (Fig. 4A), suggesting that transient PolA1 inhibition shifts early reprogramming cells toward a pro-survival state^31,32^. To validate these transcriptional changes, we examined p53 expression by immunofluorescence during early reprogramming (Fig. 4C). PolA1i-treated cells exhibited reduced p53 signal at day 3, followed by a recovery toward control levels by days 6 and 9 (Fig. 4D), indicating that PolA1 inhibition transiently attenuates p53 accumulation during the early phase of reprogramming. Because persistent suppression of p53 could permit damaged cells to persist and raise concerns about tumorigenic risk, the transient nature of this reduction is notable^33–35^. We next assessed whether the transient reduction in p53 was associated with increased γH2AX-marked DNA damage signaling^36^ (Fig. 4E). PolA1i-treated cultures contained fewer γH2AX-bright cells and displayed an overall reduction in γH2AX signal from days 3 to 16 of reprogramming (Fig. 4F-G). Thus, the transient attenuation of p53 was not accompanied by increased γH2AX-marked DNA damage pathway; instead, PolA1 inhibition was associated with reduced DNA damage-associated stress throughout reprogramming. These findings suggest that transient attenuation of p53 may contribute to improved cell survival under reprogramming-associated stress without sustained suppression of this key tumor suppressor. As an orthogonal measure of cell death, we performed a TUNEL (terminal deoxynucleotidyl transferase dUTP nick-end labeling) assay at day 16 (Fig. 4H). Consistent with the γH2AX results, PolA1i-treated cultures showed approximately a two-fold reduction in TUNEL-positive cells compared with controls (Fig. 4I). Together, these findings demonstrate that transient PolA1 inhibition reduces both DNA damage-associated stress and cell death during the establishment of emerging iPSC colonies.

**Figure 4.**
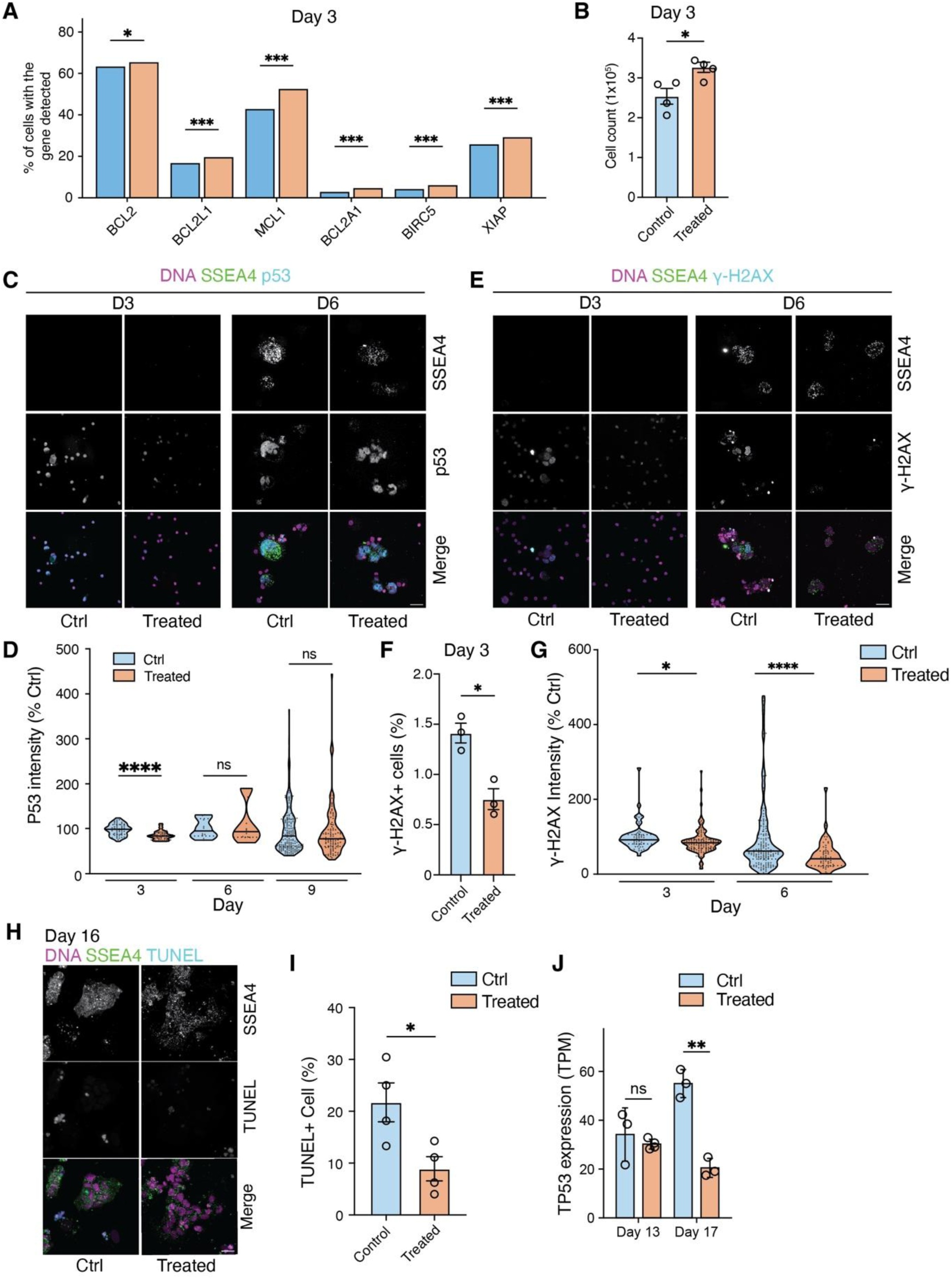
PolA1 inhibition promotes cell survival and reduces activation of cell-death pathways during reprogramming. (**A**) snRNA-seq analysis of the percentage of day 3 iPSCs during reprogramming with detectable transcripts of anti-apoptotic genes in control and PolA1 inhibitor-treated conditions, (**B**) Quantification of cell numbers on day 3 in control and PolA1 inhibitor-treated conditions. (**C**) Representative immunofluorescence images of p53 and SSEA4 during reprogramming in control and PolA1 inhibitor-treated cells and (**D**) quantification of p53 intensity from day 3 to 9. (**E**) Representative immunofluorescence images of iPSC during reprogramming on Day 3 & 6, on SSEA4 and γH2AX signals between control and PolA1i-treated condition. (**F**) Quantification of cells with bright γH2AX signal on day 3. (**G**) Quantification of basal γH2AX fluorescence intensity in control and PolA1 inhibitor-treated cells. (**H**) Representative TUNEL staining of control and PolA1 inhibitor-treated colony on day 16. (**I**) Quantification of TUNEL-positive cells in control and PolA1 inhibitor-treated colony. (**J**) TP53 transcript levels remain unchanged on day 13 but are reduced on day 17 following PolA1 inhibition. Scale bars: 25 μm (**C, E, H**).

We next asked whether reduced cellular stress translated into improved establishment of stable iPSC lines. At day 16, emerging colonies were manually picked and passaged, a procedure that imposes substantial stress on newly reprogrammed cells (Fig. 1A). Compared with control cultures, PolA1i-treated cultures generated approximately three-fold more stable iPSC lines and exhibited an approximately two-fold increase in the efficiency of stable-line establishment following passaging (Fig. S4A). Therefore, PolA1 inhibition not only increased the number of emerging colonies but also enhanced their ability to survive passaging and progress to stable iPSC lines.

Consistent with these results, transcriptome analysis of day 17 colonies revealed reduced TP53 expression together with enrichment of pathways involved in negative regulation of cell differentiation and extracellular matrix organization in PolA1i-treated colonies (Fig. 4J).

Collectively, these findings indicate that transient PolA1 inhibition improves cellular fitness during reprogramming, at least in part by reducing DNA damage-associated stress and cell death, thereby increasing the efficiency with which emerging colonies mature into stable iPSC lines.

Enhanced cellular resilience may have important implications for regenerative medicine, where substantial cell loss can occur during cell expansion, manipulation, and transplantation.

### Transient, low-dose PolA1 inhibition does not reduce DNA replication speed or affect cell cycle progression

Given the central role of PolA1 in DNA replication, we asked whether the enhanced reprogramming induced by PolA1 inhibition involves a replication-associated mechanism. From the sn multiomic data analysis, characterization of cell-cycle states revealed a higher percentage of cycling cells in Treated group than in Controls (Fig. 5A). To determine whether cells actively undergoing DNA replication at the onset of reprogramming preferentially contribute to iPSC colony formation, we performed an EdU pulse-chase tracking assay. Cells were pulsed with EdU for two hours immediately before Sendai virus transduction and then chased in EdU-free medium for five days to track their subsequent contribution to iPSC colonies (Fig. 5B). To determine whether the 2-hour pulse captured most replication-active cells, we compared it with a 48-hour EdU pulse (Fig. S5A). Extending the labeling period did not further increase the proportion of EdU^+^ cells, which remained at approximately 3%, although EdU fluorescence intensity increased substantially, consistent with greater cumulative EdU incorporation (Fig. S5B). These results indicate that a 2-hour pulse was sufficient to label most cells undergoing replication before reprogramming induction.

**Figure 5.**
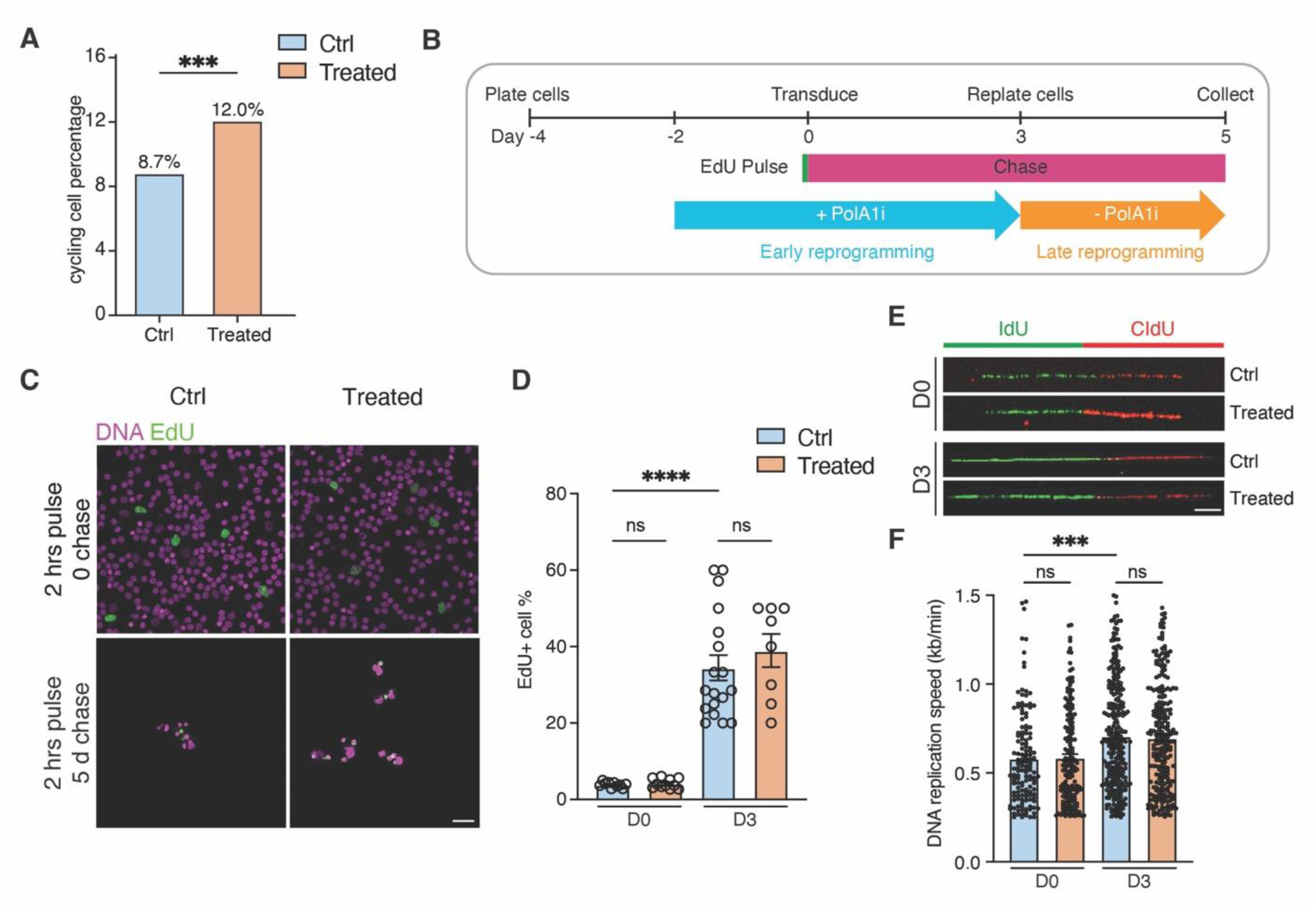
Transient, low-dose PolA1 inhibition does not reduce DNA replication speed or affect cell cycle progression. **A**) Percentage of cycling cells estimated from scRNAseq data. Seurat cell-cycle scores were calculated for each cell. Cells with S score or G2/M score > 0.1 were considered cycling cells. n = 5,992 (control), 4,662 (Treated) nuclei. (**B**) Schematic overview of the EdU pulse-chase tracking assay during the early phase of iPSC reprogramming. Representative immunofluorescence images (**C**) and quantification of the EdU-positive cell fraction (**D**) following the initial EdU pulse or the 5-day chase under control and PolA1i-treated conditions. Representative DNA fiber images (**E**) and quantification of replication-fork speed (**F**) under control and PolA1i-treated conditions on days 0 and 3. Scale bars: 25μm (**C**), 5 μm (**E**).

During the subsequent chase, the proportion of EdU^+^ cells increased from approximately 5% at day 0 to 35% by day 5 (Fig. 5C-D). Thus, cells that were undergoing DNA replication at the onset of reprogramming became progressively enriched during the early reprogramming period, suggesting that replicative cells preferentially contribute to the emerging reprogramming population.

We next focused our mechanistic studies on two biologically distinct time points: day 0 and day 3. Day 0 captures the effects of 2-day PolA1i treatment immediately before the onset of Yamanaka factor induction (pre-treatment), whereas day 3 represents a critical early reprogramming window during which Yamanaka factors drive extensive chromatin remodeling and initiate cell-fate conversion^37–39^ in the presence of PolA1i (post-treatment). Examining these two time points allowed us to distinguish molecular changes established before reprogramming from those that persist or emerge during early cell-fate transition.

Because PolA1i is known to directly target lagging-strand DNA Pola and reduce its DNA binding affinity and primer elongation^14,15^, we next asked whether the low-dose PolA1i used here affects replication-fork progression. Single-molecule DNA fiber analysis at both day 0 and day 3 revealed no significant reduction in replication-fork speed following PolA1i treatment (Fig. 5E-F). Fork speed increased from day 0 to day 3 in both control and PolA1i-treated cells (Fig. 5F), coinciding with the transition from largely quiescent PBMCs to actively cycling, reprogramming cells and suggesting the adaptation of DNA replication dynamics during early cell-fate change^40^. Together with the unchanged S-phase index (Fig. S1B), these findings indicate that 10 nM PolA1 inhibition does not measurably impair global DNA replication. Thus, the enhanced reprogramming induced by PolA1i is unlikely to result from a generalized slowdown of DNA replication.

### Transient, low dose PolA1 inhibition alters replication-coupled histone incorporation

Having found that 10 nM PolA1i inhibitor does not measurably impair global DNA replication, we next asked whether it alters chromatin state at early reprogramming time points on day 0 and day 3. The sn multiomic analysis of cycling cells showed a higher promoter fraction in PolA1i-treated cells than in controls at both time points (Fig. 6A). To directly examine chromatin accessibility in individual cells, we used NicE-view to quantify the global chromatin openness level^41^. NicE-view signal, normalized to DNA content, was higher in PolA1i-treated cells than in controls at both day 0 and day 3, with a larger increase at day 3 (Fig. 6B-C). Levels of pan-acetylated H3 (Pan-ac H3) were similarly increased (Fig. 6B-C). These differences were already detectable at day 0, before delivery of the reprogramming factors, suggesting that PolA1 inhibition alters chromatin state independently of the reprogramming program itself.

**Figure 6.**
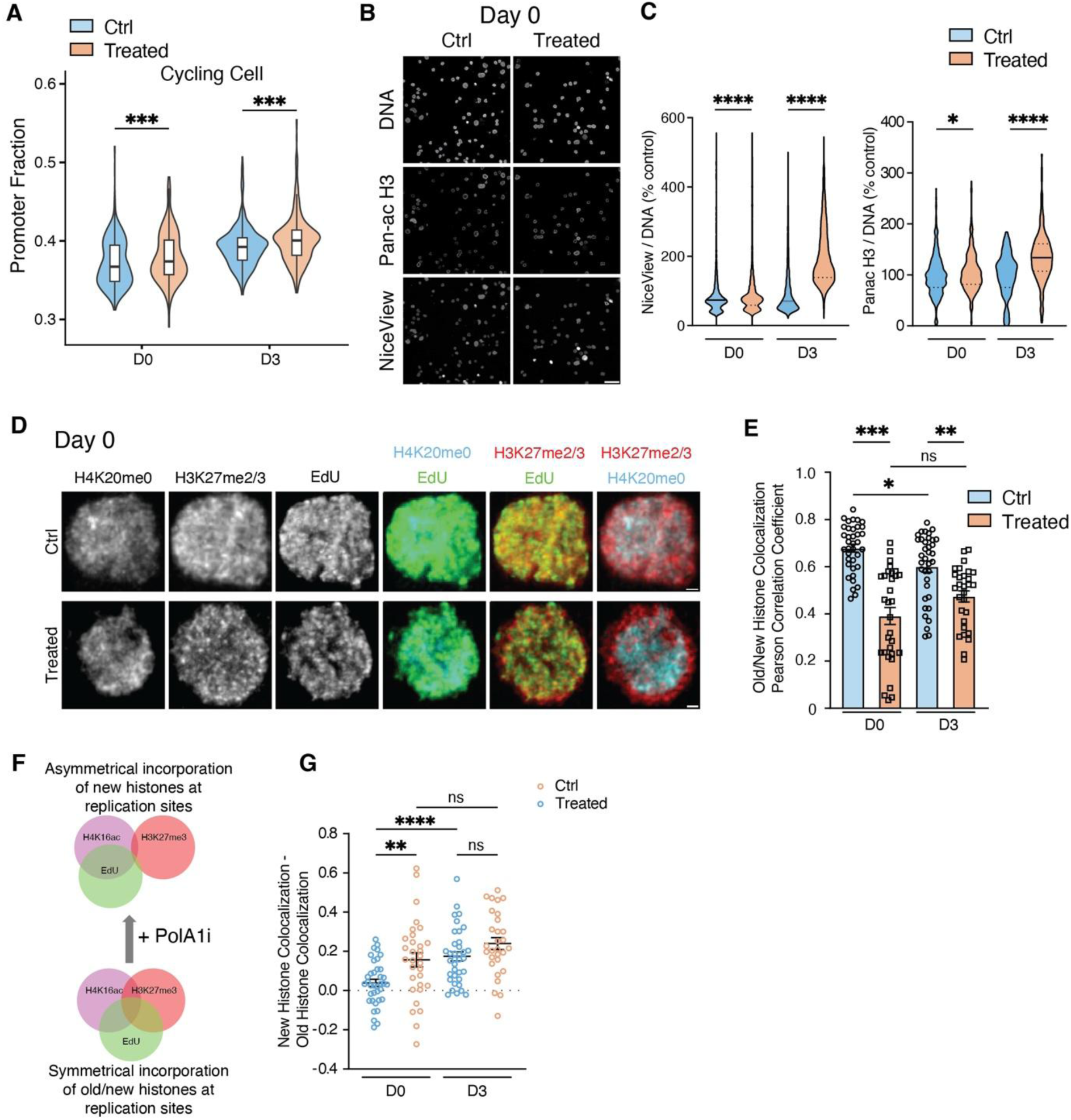
Transient, low dose PolA1 inhibition alters replication-coupled histone incorporation. (**A**) snATAC-seq analysis of the promoter fraction quantification in cycling cells from Control and Treated conditions on days 0 and 3. (**B**) Representative immunofluorescence images of DNA, pan-acetyl H3, and NiceView signals in Control and Treated PBMCs on day 0. (**C**) Quantification of DNA-normalized NiceView (left) and pan-acetyl H3 (right) intensity, Representative immunofluorescence images (**D**) and quantification of colocalization scores (**E**) between H4K20me0 (New histone) and H3K27me2/3 (Old histone) under control and PolA1i-treated conditions on days 0 and 3. (**F**) Schematic overview of colocalization across new histone, old histone, and EdU signal in active DNA replication region. (**G**) Quantification of normalized colocalization score between EdU and H4K20me0 or between EdU and H3K27me2/3 under control and PolA1i-treated conditions on day 0 and 3. Scale bars: 10 μm (**B**), 1 μm (**D**).

Since the propagation of epigenetic information is coupled to DNA replication, changes in replication dynamics could influence the inheritance of histones and their post-translational modifications (PTMs) on newly replicated DNA^42^. Previous work in the *Drosophila* male germline showed that Pola levels regulate replication-coupled incorporation of parental (old) and newly synthesized histones, with reduced Polα promoting a stem-cell-like histone incorporation pattern^12^. We therefore asked whether 10 nM PolA1 inhibition similarly alters replication-coupled histone incorporation during human PBMC reprogramming.

To distinguish old from new histones, we used the well-characterized old-histone-enriched mark H3K27me2/3^43,44^ and the new-histone-enriched mark H4K20me0^45^ to examine their localization relative to nascent DNA in S-phase PBMCs pulse-labeled with EdU (Fig. 6D). As expected, EdU-negative, unreplicated regions were enriched for H3K27me2/3 but depleted of H4K20me0 (Fig. 6D), consistent with their respective enrichment on old and newly synthesized histones.

Using stringent conditions to remove non-chromatin-bound histones and high-resolution imaging (Methods), we found that H3K27me2/3 and H4K20me0 exhibited significantly greater spatial separation in PolA1i-treated PBMCs than in control cells at S phase following two days of PolA1i treatment before reprogramming induction (day 0, Fig. 6E). This pattern is consistent with a shift from symmetric incorporation of old and new histones at replication sites toward a more asymmetric incorporation of new histones (Fig. 6F). To quantify this shift, we calculated the difference between new histone (H4K20me0) and old histone (H3K27me2/3) colocalization with EdU (Fig. 6G). Notably, this difference was significantly higher in Treated cells than in Control cells at day 0, indicating a higher association between new histones and nascent DNA than old histones following PolA1i treatment. These findings indicate that 10 nM PolA1 inhibition establishes an altered replication-coupled histone incorporation pattern before the initiation of reprogramming.

We next examined whether this pattern was maintained following reprogramming induction. After 3 days of continued PolA1i exposure following viral delivery of the reprogramming factors, H3K27me/3 and H4K20me0 remained significantly more separated in Treated cells than in Control cells (day 3, Fig. 6E). Importantly, the day 3 pattern should be considered in the context of the additional changes induced by the reprogramming program itself. In control cells, continued exposure to the reprogramming factors was associated with greater spatial separation between H3K27me2/3 and H4K20me0, as well as between H3K27me2/3 and EdU, whereas H4K20me0 showed increased colocalization with EdU at day 3 compared with day 0 (Fig. 6G). These changes are consistent with a progressive shift in histone incorporation toward a pattern associated with a more stem-cell-like state during reprogramming. In contrast, continued PolA1i exposure produced no significant further changes in these colocalization patterns in PolA1i-pretreated cells (Fig. 6E, G). Thus, whereas control cells undergo additional changes in replication-coupled histone incorporation following reprogramming induction, PolA1i pre-treated cells, which already exhibited an altered pattern before reprogramming, show relatively little further change in this feature.

Together, these findings suggest that 10 nM Pola inhibition selectively alters replication-coupled histone incorporation without substantially altering global replication-fork progression. The altered pattern established before reprogramming is reminiscent of the histone incorporation pattern associated with reduced Polα levels in the Drosophila male germline. The limited additional change following reprogramming induction in PolA1i-pretreated cells raises the possibility that PolA1 inhibition places somatic cells in a chromatin state that is more permissive or poised for subsequent acquisition of a stem-cell-like state.

### Transient, low-dose PolA1 inhibition enhances human PBMC reprogramming in a two-step treatment regimen

In our experiments, PolA1 inhibitor (PolA1i) treatment spanned 5 days: 2 days of pre-treatment before Sendai-virus transduction at Day 0, and 3 days of post-treatment afterwards (Fig. 7A). In addition to the full treatment (Pre + Post, PP), we collected samples treated only before (Pre) or only after (Post) transduction, together with vehicle controls (Ctrl), for 10x multiome profiling at Days 0, 3 and 7 (Fig. 7A). In total, 10 libraries were analyzed. Adding the four new libraries (D3 Pre, D3 Post, D7 Pre and D7 Post) to the original six did not change the overall structure of the joint UMAP, which resolves 15 cell types (Fig. 7B; Fig. S7A-B). Although most cells still retain clear PBMC features, three of these cell types (13, 14 and 15) are associated with reprogramming (Fig. 7B–D).

**Figure 7.**
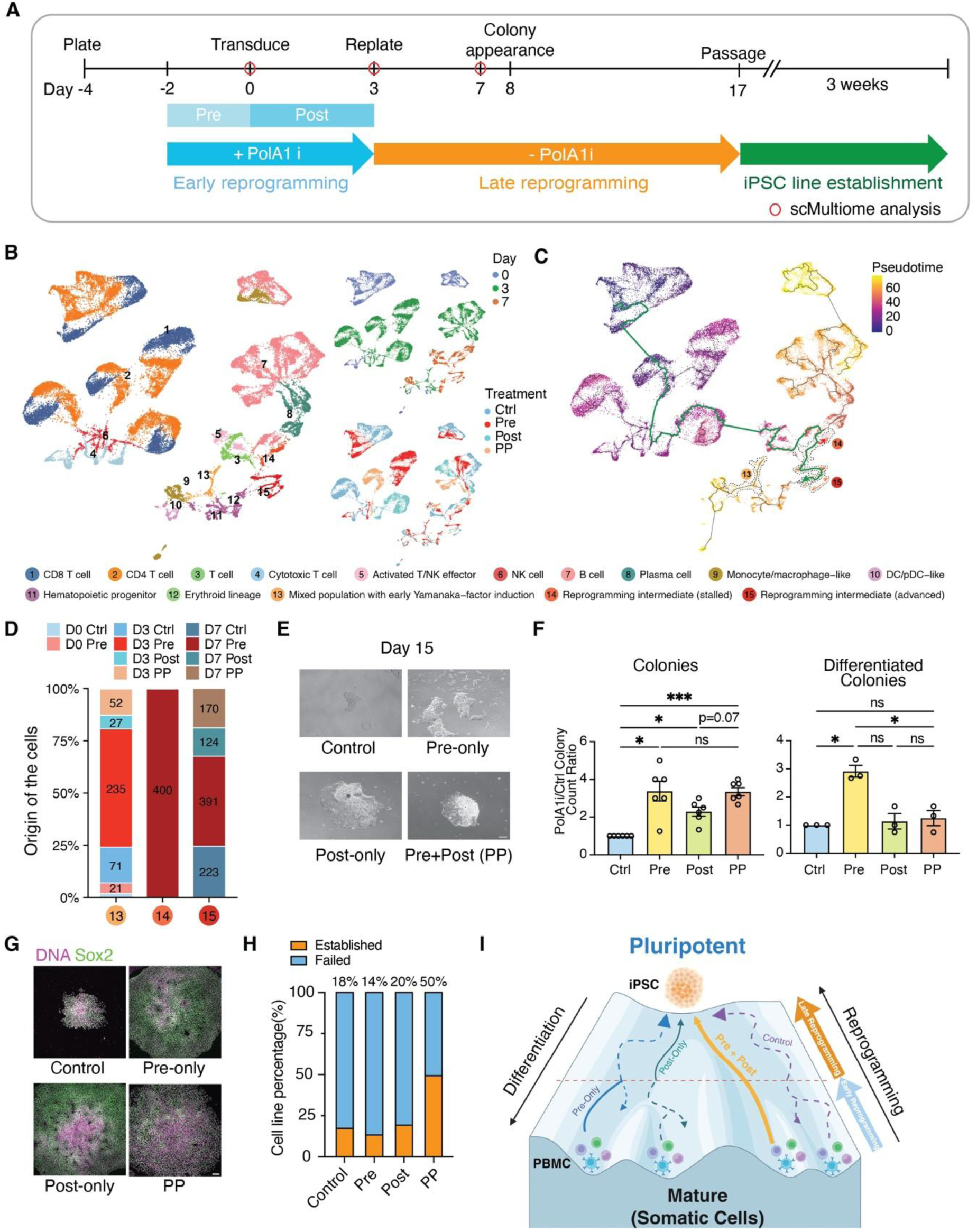
Transient, low-dose PolA1 inhibition primes human PBMCs for sustained gains in reprogramming efficiency with pre- and post-treatment. (**A**) Schematic overview of the PBMC-to-iPSC reprogramming protocol with transient PolA1 inhibition Pre- and Post-treatment. (**B**) UMAP of integrated snRNA-seq and snATAC-seq data from all 10 libraries. (**C**) Monocle3 pseudotime trajectory on the same embedding, colored by pseudotime. The green line highlights the trajectory from the starting population to the advanced reprogramming intermediates (cell type 15), and dashed outlines enclose cell types 13, 14 and 15. The small red arrow marks where cells exit the reprogramming trajectory. (**D**) Origin of the cells in the three reprogramming-associated cell types (13, 14 and 15). Representative brightfield images (**E**) of iPSC colonies and quantification of colony count ratios (**F**) on day 15 under control, Pre-, Post-, Pre + Post (PP) treatment conditions. (**G**) Representative immunofluorescence images of SOX2^+^ colonies with control and alternative PolA1i treatment conditions on day 15. (**H**) Quantification of established and failed cell lines percentage across control and alternative PolA1i treatment conditions, n = 16 (Ctrl), 22 (Pre), 15 (Post), 24 (PP). (**I**) Graphical model of the proposed effect of PolA1i treatment timing on human PBMC reprogramming process. Scale bars: 100 μm (**E**, **G**).

Cell type 13 is a mixed population that contains cells from multiple days and treatments and expresses low levels of the Yamanaka factors (POU5F1, SOX2 and KLF4) alongside PBMC markers (Fig. S7A). Within cell type 13, cells from D3 Pre were the most abundant (Fig. 7D). This suggests that pre-treatment promotes entry into the earliest stage of iPSC reprogramming most when most cells are still carrying PBMC identity (Fig. S7D).

Cell type 15 represents the most advanced iPSC intermediates, with the highest expression of early iPSC genes (e.g., POU5F1, SOX2, KLF4, LIN28B and DPPA4; Fig. S7A) in 92.8% of cells (Fig. S7D). All cells of cell type 15 came from Day 7, from all four conditions (Fig. 7D). We drew a trajectory from the starting population to cell type 15 to illustrate the reprogramming route (green line, Fig. 7C), and 87.4% of cell type 15 cells lay on it (Fig. S7C). In contrast, cell type 13 did not lie on this trajectory (Fig. S7C). Because it retains PBMC gene expression, this mixed population may have been pulled away from the reprogramming route when the UMAP was constructed.

Cell type 14 came almost entirely from D7 Pre (Fig. 7D), and only 40.4% of its cells (162 of 401) were on the reprogramming trajectory (Fig. S7C). The remaining 239 cells (59.6%) exited the trajectory before reaching the advanced intermediates (red arrow, Fig. 7C). Compared with cell type 15, these cells express pluripotency genes at lower levels and more often retain lineage genes (Fig. S7A, D). Cell types 14 and 15 together made up 39.2% of D7 Pre cells, more than any other D7 group (Fig. S7B). Together with the enrichment of D3 Pre cells in cell type 13, our results suggest that pre-treatment may promote reprogramming at the early stage.

However, without post-treatment, many cells fail to complete the process and stall (Fig. 7C). To further confirm how different treatments affect the reprogramming process, we collected samples at a much later time point, Day 15, to quantify the reprogramming efficiency in different groups. The Pre-treatment group indeed had more colonies, but many of them were already differentiated (Fig. 7E), consistent with a large share of Pre cells being stalled in the multiome data and later differentiating (Fig. 7C).

Quantification of colony numbers on day 15 supported this observation (Fig. 7F). Compared with controls, all three treatment regiments increased the total number of colonies: Pre-only by 3.4-fold, Post-only by about 2.3-fold, and Pre + Post (PP) by about 3.3-fold (Fig. 7F, left). The pre and PP groups yielded similar numbers of colonies, while the Post-only group tended to show a smaller increase than PP (*P*= 0.07). The colonies themselves, however, differed in quality. Differentiated colonies rose about 2.9-fold in the Pre-only group, but did not change significantly in the Post-only or PP groups (Fig. 7F, right).

We then assessed the pluripotent state of the day 15 colonies by SOX2 immunostaining (Fig. 7G). Qualitatively, all of the PolA1i-treated groups showed an overall larger colony morphology comparing with the control group. While colonies in all groups showed SOX2 signal demonstrating their pluripotency, the PP group showed the most compact and uniform SOX2 signal across its entire colony area, suggesting a higher quality of colony formed near the end of the reprogramming stage on day 15.

Finally, we tested whether these colonies could be established as stable iPSC lines after manual picking and passaging on day 17 (Fig. 7A). About half of the colonies picked from the PP condition gave rise to established lines, compared with roughly 14-20% in other conditions (Fig. 7H). Thus, although Pre-only treatment increased colony number, it did not improve line establishment, and Post-only treatment was likewise insufficient. Taken all together with the multiomic data and the results, we propose that pre-treatment promotes the earliest stage of reprogramming, but the continued PolA1i inhibition after Yamanaka factors delivery is needed for human PBMCs to progress through reprogramming and remain undifferentiated, so that combined Pre + Post treatment is required to yield the most stable iPSC lines (Fig. 7I).

## Discussion

Our study demonstrates that transient, partial inhibition of the Polα catalytic subunit PolA1 accelerates and enhances the reprogramming of human PBMCs into iPSCs.

Mechanistically, transient PolA1 inhibition promotes early chromatin and transcriptional remodeling, characterized by increased chromatin accessibility, more rapid progression toward a pluripotent transcriptional state, and coordinated changes in stress-response and cell survival pathways consistent with improved cellular fitness during reprogramming. Beyond increasing iPSC colony formation, PolA1 inhibition promotes the establishment of colonies with higher expression of pluripotency markers and enhanced differentiation potential. Together, these findings provide proof-of-concept that controlled modulation of the DNA replication machinery can enhance somatic cell reprogramming and identify replication machinery as an underappreciated regulator of epigenetic plasticity and cell-fate transitions.

### The magnitude and timing of DNA replication modulation determine its effects on cellular plasticity

Our findings suggest that the magnitude of DNA replication modulation is a key determinant of whether perturbing the replication machinery enhances cellular plasticity or instead compromises normal cell function. This concept may help reconcile our findings with previous studies showing that disruption of replication-coupled chromatin assembly can impair stem cell differentiation and development.

DNA replication machinery is increasingly recognized as playing dual roles in genome duplication and epigenetic inheritance through replication-coupled parental histone recycling^46–49^. Among these factors, Mcm2, Ctf4, and Polα form a histone-transfer axis that coordinates parental histone recycling to the lagging strand^11,50,51^. Consistent with the importance of this process, separation-of-function mutations that disrupt histone chaperone activity while preserving core helicase or polymerase functions, including *mcm2-2A* and *pole3*, produce pronounced asymmetry in parental histone inheritance and impair embryonic stem cell differentiation and embryonic development^50–55^.

Our approach differs fundamentally from these genetic perturbations. Rather than disrupting histone-transfer activity directly, we modestly reduced Polα dosage through heterozygosity^13^ or transiently inhibited PolA1 activity using an optimized low-dose treatment. We propose that this level of perturbation is sufficient to alter replication-coupled chromatin inheritance and increase chromatin plasticity without inducing the broad disruption of chromatin organization associated with more severe perturbations.

Interestingly, previous studies have shown that more substantial perturbation of DNA replication progression during early mouse embryogenesis can promote cellular reprogramming toward totipotency^56^. In contrast, our optimized conditions produced no detectable changes in S-phase progression across multiple experimental systems^12,13^, and DNA fiber analyses showed minimal effects on replication fork progression speed. These findings suggest that altered replication speed itself is unlikely to account for the enhanced cellular plasticity observed here. Instead, our results point to a distinction between perturbing the mechanics of DNA replication and modulating the chromatin consequences of replication.

Together, these observations support a model in which the magnitude of DNA replication perturbation influences its biological outcome. Whereas severe disruption of replication-coupled chromatin assembly can compromise epigenetic inheritance and normal development^53^, modest and transient Polα modulation may create a permissive window in which chromatin inheritance becomes sufficiently plastic to facilitate cell-fate transitions while preserving overall replication progression and cellular fitness. We propose that this balance may represent a general principle by which controlled modulation of DNA replication machinery can enhance cellular plasticity without incurring the detrimental consequences associated with more extensive replication stress.

The use of PolA1 inhibition during iPSC reprogramming also provides an opportunity to examine the temporal requirements for replication modulation during cell-fate transitions. We found that transient pre-treatment with a low, non-disruptive dose of PolA1 inhibitor enhanced cellular stress resistance and improved subsequent reprogramming efficiency even before induction of the reprogramming factors. Continued PolA1 inhibition following reprogramming induction was also required to fully realize these benefits, suggesting that the timing of replication modulation is an important determinant of its effects. One possibility is that early PolA1 inhibition establishes a permissive or “primed” cellular state, whereas continued modulation reinforces or sustains the chromatin and transcriptional remodeling associated with acquisition of pluripotency (Fig. 7B-C). Although the molecular basis of these temporally distinct effects remains to be determined, these findings suggest that both the magnitude and duration of replication perturbation are important parameters for manipulating cellular plasticity.

### Replication modulation as a potential conserved regulator of cellular plasticity

Cellular reprogramming and regenerative medicine have transformed our ability to manipulate cell fate, creating new opportunities to restore tissues damaged by aging, injury, and disease. A central challenge, however, is to identify mechanisms that can increase cellular plasticity while preserving genome integrity and normal tissue function.

Our findings, together with previous observations *in vivo*^12,13^, suggest that the DNA replication program itself may serve as a regulatory interface between DNA replication, chromatin inheritance, and cellular plasticity. By modulating the activity of the Polα complex, an essential initiator of lagging-strand DNA synthesis, we found that controlled perturbation of replication machinery establishes a permissive state for cell-fate transitions. In vivo, modest Polα modulation enables differentiated cells to undergo dedifferentiation and compensate for stem cell loss during aging and tissue damage while retaining the capacity to redifferentiate into functional cell types^13^. *Ex vivo*, transient pharmacological inhibition of PolA1 enhances iPSC generation from human PBMCs. Together, these findings suggest that cellular plasticity in distinct biological contexts may be sensitive to the activity of the same core replication machinery, raising the possibility that controlled modulation of replication-coupled chromatin inheritance represents a broadly applicable mechanism for regulating cell-state transitions.

### Translational implications of controlled replication modulation

The ability to manipulate cellular plasticity without permanently altering the genome has important implications for regenerative medicine and cell-based therapies. Our findings provide proof-of-concept that transient modulation of the DNA replication machinery can improve both the efficiency and robustness of somatic cell reprogramming. PolA1 inhibition increased the formation of iPSC colonies, promoted more rapid progression toward a pluripotent transcriptional state, and increased the establishment of stable iPSC lines. Importantly, PolA1i-treated cells also displayed enhanced recovery following exposure to diverse external stresses, suggesting that the benefits of replication modulation extend beyond increasing the initial efficiency of reprogramming to improving cellular resilience during cell-state transitions.

These properties may be particularly advantageous for applications in which reprogramming efficiency and cellular survival represent major barriers to generating high-quality cell products^57^. Because PolA1 inhibition is transient and does not require permanent genetic modification of the replication machinery, this approach may offer a potentially controllable means of manipulating cell-state plasticity. The observation that continued PolA1 inhibition after reprogramming-factor induction further enhanced the phenotype also suggests that the timing and duration of replication modulation can be optimized according to the specific stage of cell-state transition. More broadly, the principle established here may be applicable not only to iPSC generation but also to other contexts in which controlled increases in cellular plasticity could improve regeneration or cell-state conversion.

Several considerations will be important for translating this strategy. First, the magnitude and duration of PolA1 inhibition must be carefully controlled. Our results indicate that modest, transient modulation can enhance cellular plasticity without detectable disruption of replication progression, whereas more extensive perturbation of replication or replication-coupled chromatin assembly can compromise cellular function. Defining the dose and temporal window that maximize plasticity while minimizing replication stress will therefore be essential. Second, although our analyses did not reveal increased DNA damage or persistent p53 suppression, and instead showed reduced DNA damage-associated signaling and cell death during reprogramming, longer-term genomic stability and tumorigenic potential will require direct evaluation in future studies. Third, it will be important to determine whether the effects of PolA1 modulation are conserved across different starting cell types and whether the approach can be applied selectively to cells undergoing desired cell-state transitions without affecting neighboring or differentiated cells.

An additional consideration is whether transient replication modulation can be integrated with existing reprogramming or regenerative strategies to reduce the intensity or duration of other interventions. The ability of PolA1 inhibition to establish a more permissive cellular state before reprogramming-factor induction raises the possibility that replication modulation could be used as a preparative step, while continued short-term modulation may support subsequent chromatin and transcriptional remodeling. Such strategies could potentially improve the consistency and scalability of cell-state conversion while reducing cellular loss during stressful transition periods.

Overall, our findings identify DNA replication machinery as an experimentally tractable regulator of cellular plasticity. Rather than broadly disrupting DNA replication, precisely calibrated and transient modulation of Polα provides a means to alter the chromatin consequences of replication while preserving overall cellular fitness. This work establishes a framework for exploring replication modulation as a strategy to improve cellular reprogramming and, more broadly, to control cell-state transitions for regenerative and therapeutic applications.

## Supplementary figures and figure legends

**Figure S1:**
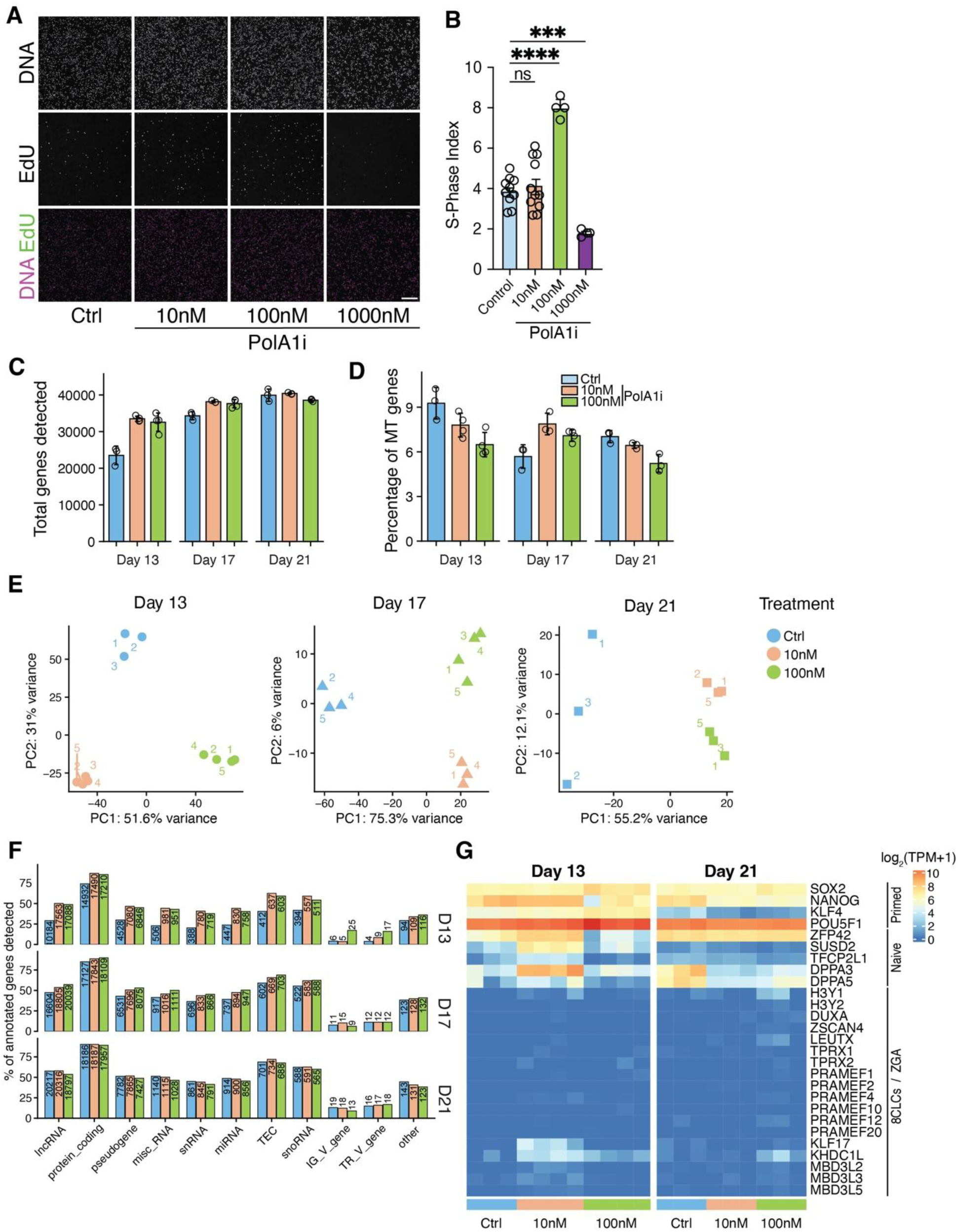
Low-dose PolA1 inhibition does not affect S-phase progression. (**A**) Representative images of EdU-positive cells treated with vehicle control or different doses of PolA1 inhibitor. Scale bar, 10 µm. (**B**) Quantification of the S-phase index under the indicated treatment conditions.

**Figure S2:**
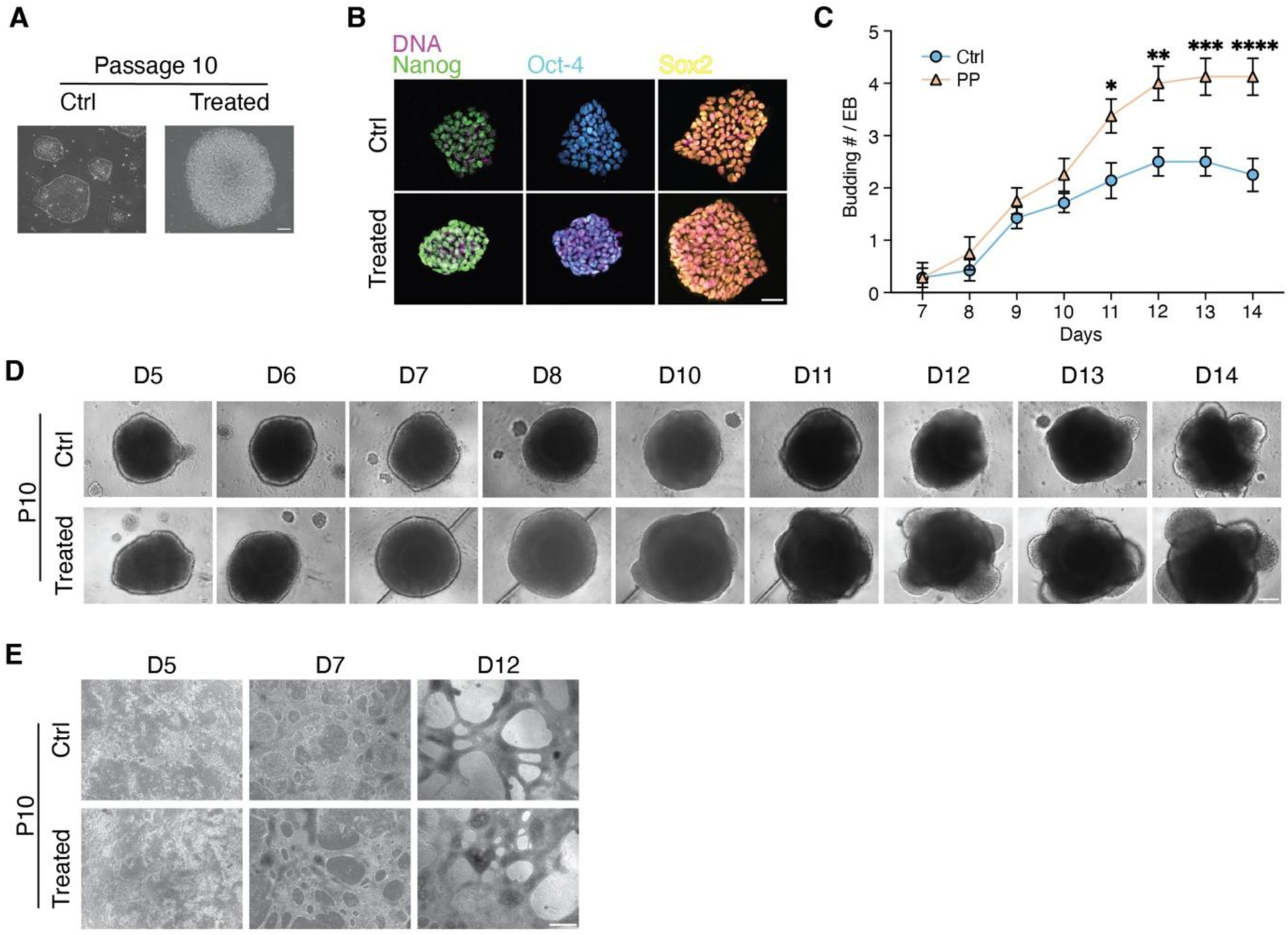
Later passage stabilized PolA1i-derived iPSC lines maintains pluripotency and enhanced differentiation advantage. (**A**) Representative bright-field images of late-passage (P10) iPSC lines derived from control- or PolA1i-treated cells. (**B**) Representative immunofluorescence images showing expression of the core pluripotency factors SOX2, NANOG, and OCT4 in control- and PolA1i-derived iPSC late passage (P10) lines. (**C**) Quantification of number of buds per EB from day 7-14 differentiated from late passage (P10) control and PolA1i-derived iPSC. (**D**) Representative bright-field images of late passage-10 iPSC-derived EBs with buds during differentiation from day 5 to day 14. (**E**) Representative bright-field images of cardiomyocyte differentiated from late passage iPSC (P10) on days 5, 7, and 12. Scale bars: 100 μm (**A**), 50 μm (**B**), 200 μm (**D**), 500 μm (**E**).

**Figure S3.**
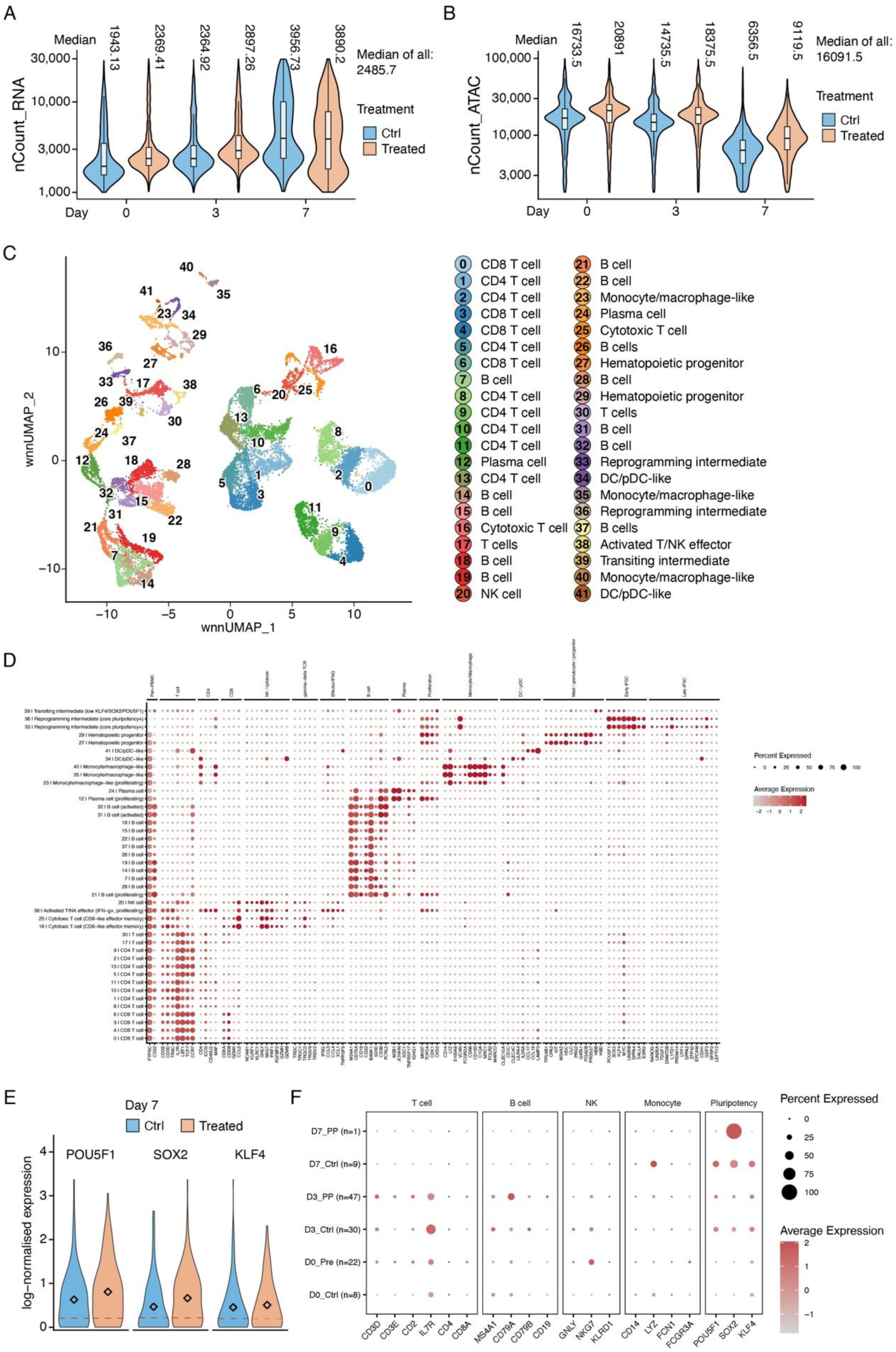
Quality control, cluster annotation, and marker expression in the multiomic dataset. (**A, B**) Distribution of RNA counts (nCount_RNA; A) and ATAC counts (nCount_ATAC; B) per nucleus in control and PolA1i-treated samples at days 0, 3, and 7, after quality filtering. Values above each violin are the sample medians (the median of all nuclei is shown on the right of each panel). Box plots indicate the median and interquartile range. (**C**) WNN UMAP showing all 42 clusters (0–41) with the cell type assigned to each cluster. (**D**) Dot plot of marker gene expression across clusters 0–41. Marker genes are grouped by category. Dot size shows the percentage of cells expressing the gene, and color shows the scaled average expression. (**E**) Log-normalized expression of POU5F1, SOX2, and KLF4 in day 7 control and PolA1i-treated cells. Diamonds mark the mean, and the red dashed line marks average expression of all genes annotated. (**F**) Dot plot of lineage and pluripotency marker expression in cluster 39 cells, split by day and treatment. The number of cells in each group is given in parentheses. Dot size shows the percentage of cells expressing the gene, and color shows the scaled average expression.

**Figure S4:**
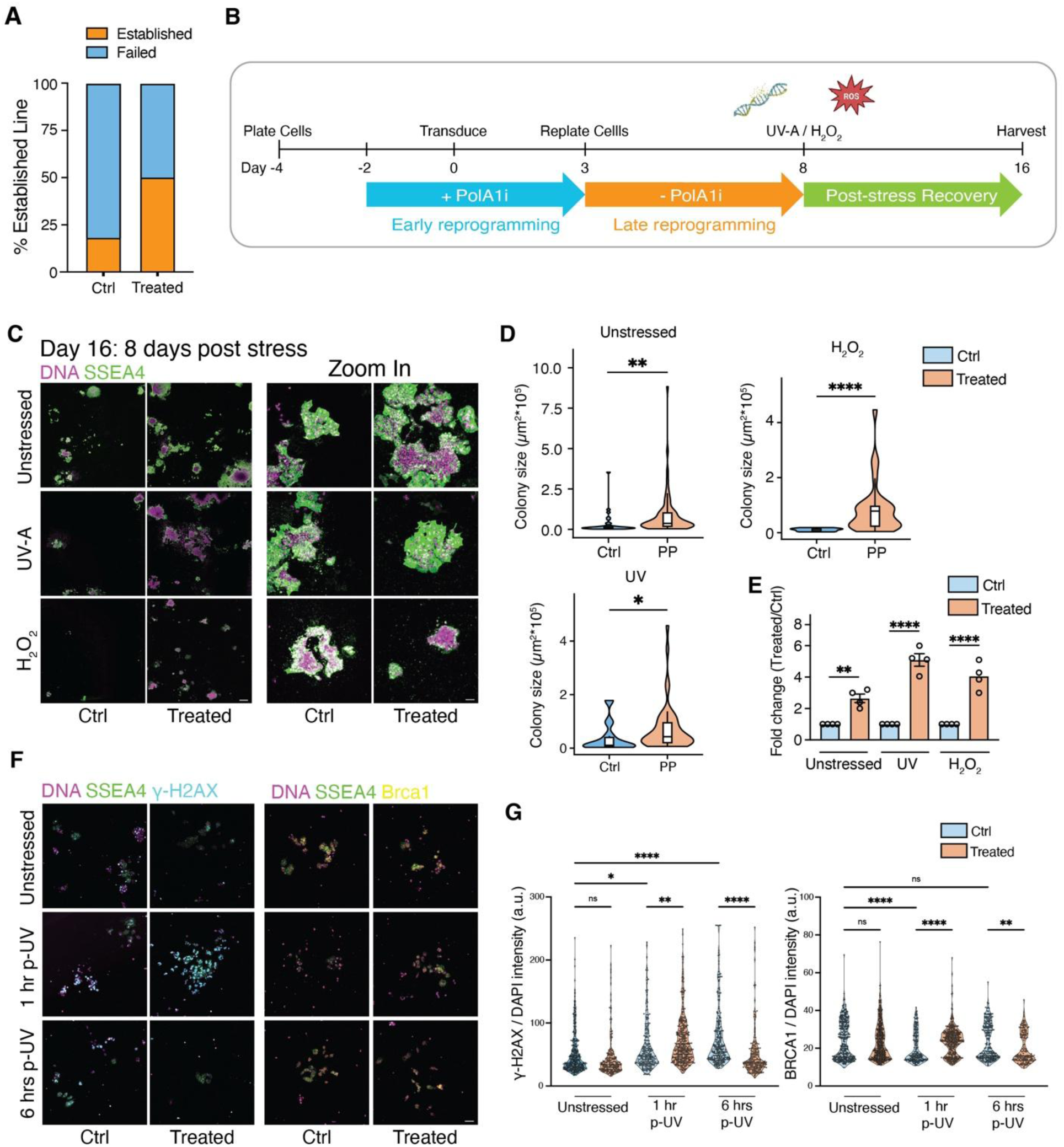
PolA1 inhibition enhances cellular resilience during reprogramming. (**A**) Percentage of iPSC lines generated from control and PolA1 inhibitor-treated conditions. (**B**) Schematic overview of the experimental workflow, with UV-A or H₂O₂ exposure on reprogramming cells at day 8. (**C**) Representative immunofluorescence images of SSEA4+colonies 8 days after stress exposure. (**D**) Quantification of colony number and size post-stress treatment conditions, and (**E**) fold enrichment of total SSEA4+ colony area in PolA1i-treated cultures relative to controls under unstressed, UV-A exposure, and H₂O₂ treatment conditions. (**F**)Representative immunofluorescence images of γH2AX and BRCA1, and (**G**) Quantification of normalized γH2AX and BRCA1 immunofluorescence intensities, before and at 1 and 6 hours post UV-A exposure. Scale bars: 500 μm (**C**), 50 μm (**C-Zoom in**) 50 μm (**F**).

**Figure S5:**
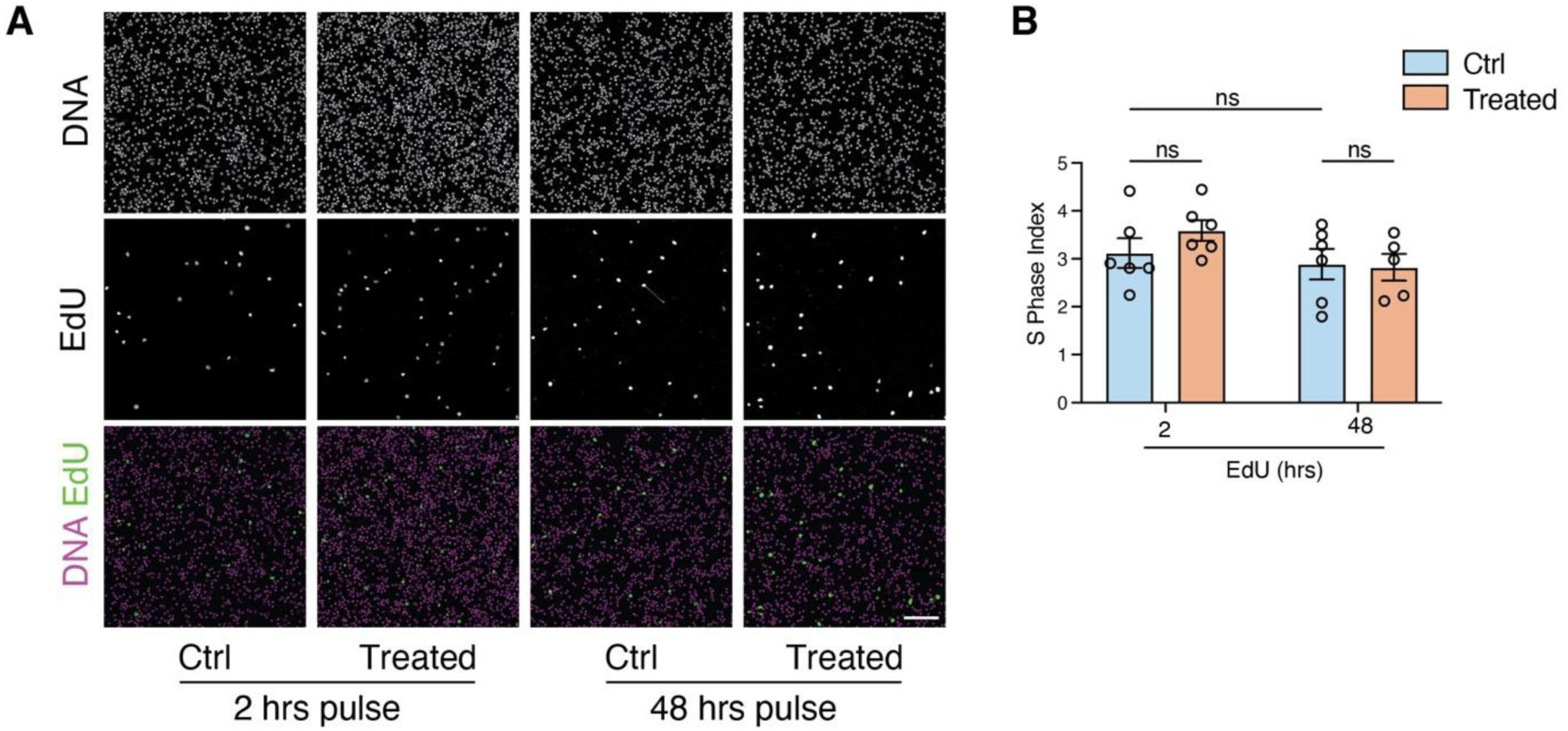
Prolonged EdU pulse does not increase labeled S-phase cells. Representative immunofluorescence images (**A**) and quantification (**B**) of the EdU-positive cell fraction with initial 48 h pulse and chase till day 5 under control and PolA1i-treated conditions. Scale bars: 25 μm (**A**).

**Figure S6:**
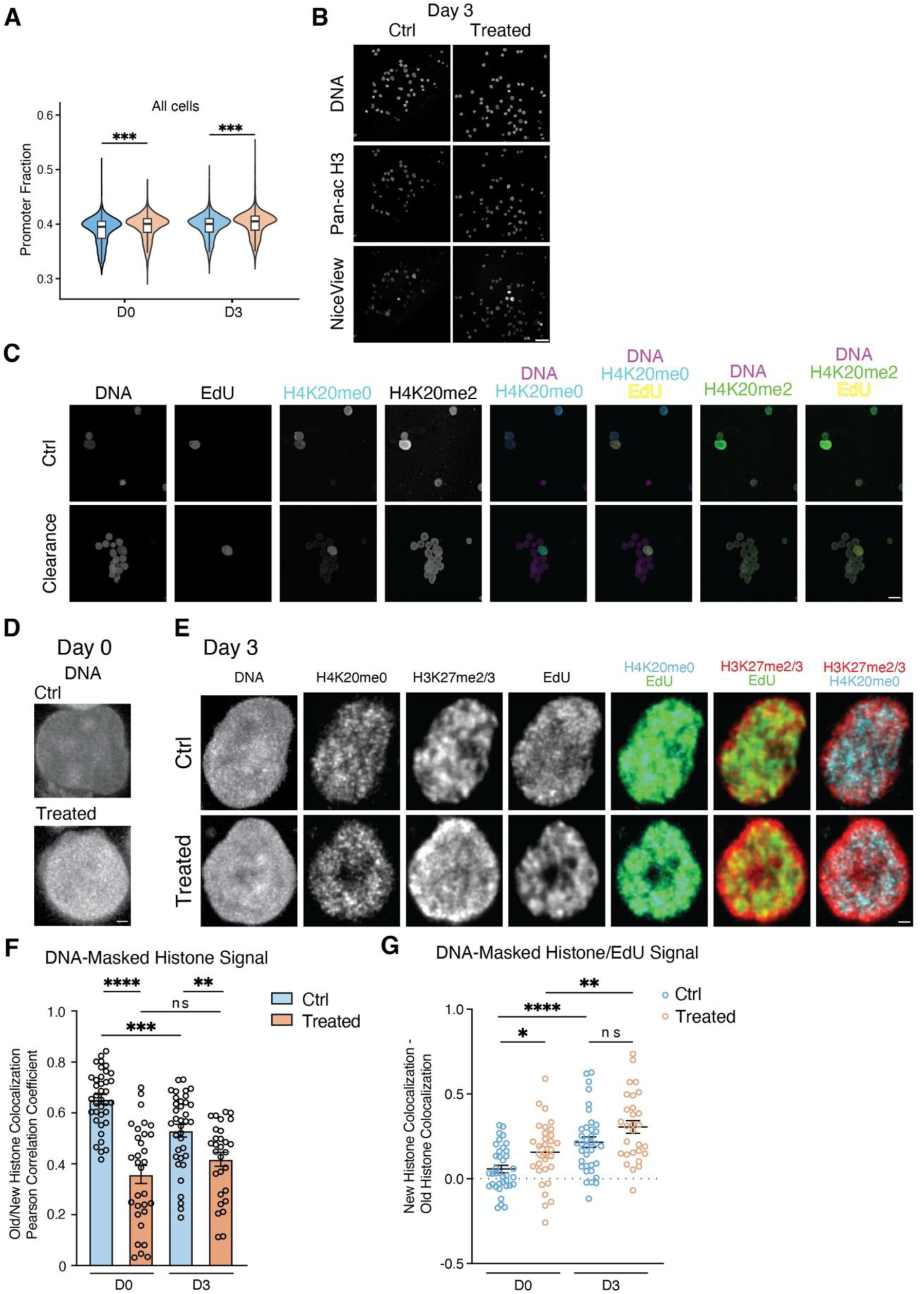
Transient, low dose PolA1 inhibition alters replication-coupled histone incorporation. (**A)** snATAC-seq analysis of the promoter fraction quantification in all cells from Control and Treated conditions on days 0 and 3. (**B**) Representative immunofluorescence images of DNA, pan-acetyl H3, and NiceView signals in Control and Treated PBMCs on day 3. (**C**) Representative immunofluorescence images of the nuclear histone signal with and without clearance buffer processing. (**D**) Representative immunofluorescence images of DNA-only channel for day 0 Control and Treated cell samples. Representative immunofluorescence images (**E**) and quantification of colocalization scores (**F**) between H4K20me0 (New histone) and H3K27me2/3 (Old histone) under control and PolA1i-treated conditions on days 0 and 3 from signals quantified with DNA signal masking in 3D space. (**G**) Quantification of normalized colocalization score between EdU and H4K20me0 or between EdU and H3K27me2/3, under control and PolA1i-treated conditions on day 0 and 3 based on signals quantified using DNA signal as masking in 3D space. Scale bars: 10 μm (**B-C**), 1 μm (**D-E**).

**Figure S7:**
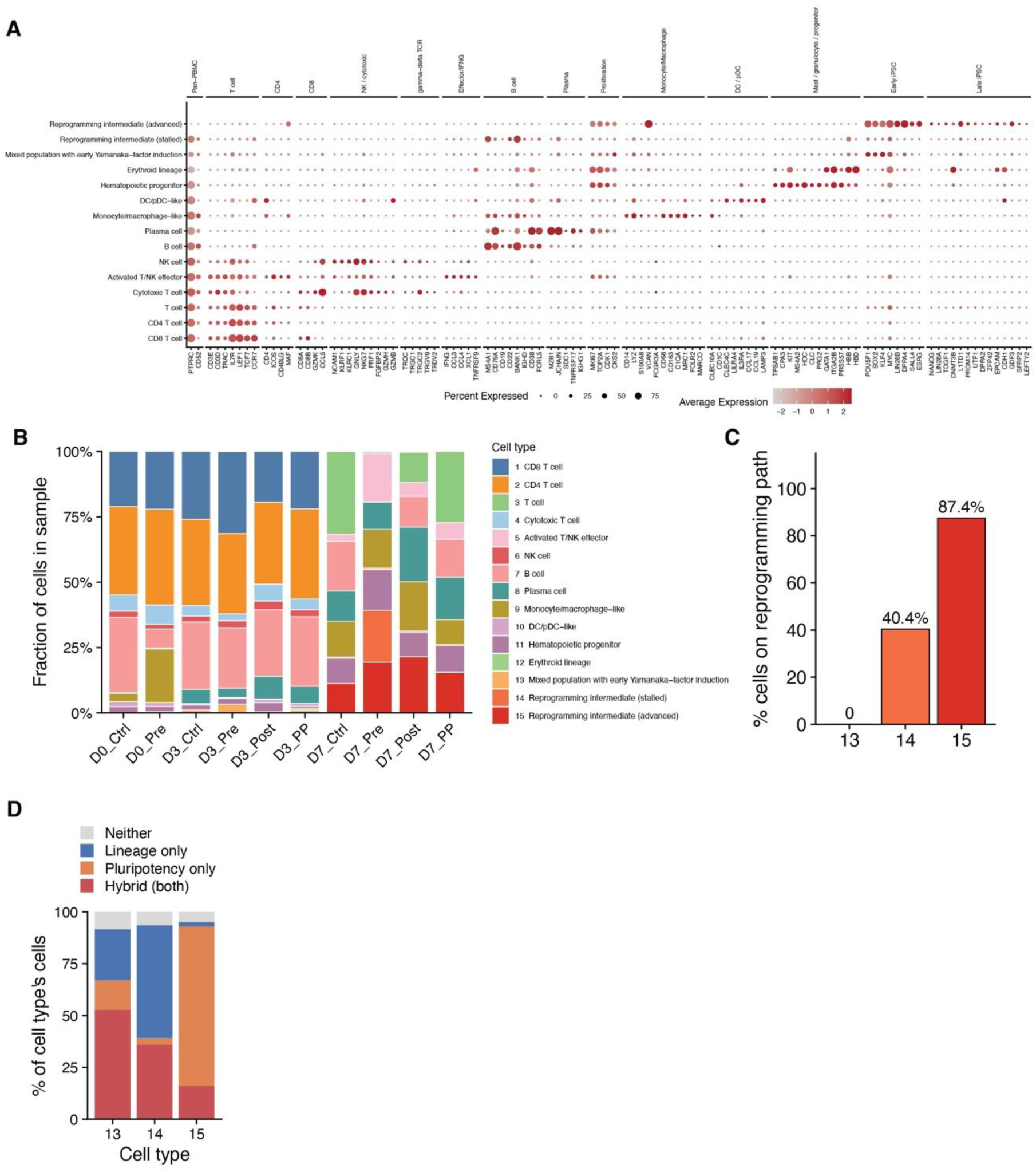
Characterization of cell types identified in scMultiome. (**A**) Dot plot of marker gene expression across the 15 cell types. Genes are grouped by category. (**B**) Fraction of cells in each of the 10 libraries assigned to each cell type. Cell type numbers and colors are as in Fig. 7B. (**C**) Percentage of cells in cell types 13, 14 and 15 that lie on the reprogramming trajectory (green line in Fig. 7C). Numbers above the bars show cells on the trajectory over total cells in the cell type. (**D**) Composition of cell types 13, 14 and 15 according to expression of PBMC lineage and pluripotency genes.

