## Supplementary material for "Transient Partial Inhibition of DNA Polymerase α Promotes Chromatin Plasticity and Enhances Human Cell Reprogramming": Materials and Methods

### Key resources table

| REAGENT or RESOURCE | SOURCE | IDENTIFIER |
| --- | --- | --- |
| **Antibodies** |  |  |
| Anti-ssDNA Antibody, clone 16-19 ZooMAb® Mouse Monoclonal | Sigma | Cat# ZMS1042; RRID: AB_3754681 |
| BD™ Purified Mouse Anti-BrdU Clone B44 (RUO (GMP)) | BD Biosciences | Cat# 347580; RRID: AB_10015219 |
| Anti-BrdU antibody [BU1/75 (ICR1)] - Proliferation Marker | Abcam | Cat# ab6326; RRID: AB_305426 |
| SSEA4 Monoclonal Antibody (eBioMC-813-70 (MC-813-70)), Alexa Fluor™ 488, eBioscience™ | Invitrogen | Cat# 53-8843-42; RRID: AB_10609335 |
| Goat polyclonal anti-SOX2 | R&D | Cat# AF2018; RRID: AB_355110 |
| Nanog (D73G4) Rabbit Monoclonal Antibody | Cell Signaling Technology | Cat# 4903; RRID: AB_10559205 |
| Purified anti-human SUSD2 Antibody | BioLegend | Cat# 327401; RRID: AB_940656 |
| Histone H3K27me3 antibody (mAb) | Active Motif | Cat# 61017; RRID: AB_2614987s22 |
| Phospho-Histone H2A.X (Ser139) (D7T2V) Mouse Monoclonal Antibody | Cell Signaling Technology | Cat# 80312; RRID: AB_2799949 |
| p53 (1C12) Mouse Monoclonal Antibody | Cell Signaling Technology | Cat# 2524; RRID: AB_331743 |
| Cardiac Troponin T Monoclonal Antibody (13-11) | Invitrogen | Cat# MA5-12960; RRID: AB_11000742 |
| alpha Actinin 2 Recombinant Rabbit Monoclonal Antibody (7H1L69) | Invitrogen | Cat# 701914; RRID: AB_2688290 |
| SOX1 Polyclonal antibody | ProteinTech | Cat# 20744-1-AP; RRID: AB_2878732 |
| Brachyury Polyclonal Antibody | Invitrogen | Cat # PA5-46984; RRID: AB_2610378 |
| SOX17 Polyclonal antibody | ProteinTech | Cat# 24903-1-AP; RRID: AB_2879789 |
| Goat Anti-Rabbit IgG H&L (Alexa Fluor® 488) | Abcam | Cat# ab150077; RRID: AB_2630356 |
| Donkey anti-Mouse IgG (H+L) Highly Cross-Adsorbed Secondary Antibody, Alexa Fluor™ 488 | Invitrogen | Cat# A-21202; RRID: AB_141607 |
| Donkey anti-Rabbit IgG (H+L) Highly Cross-Adsorbed Secondary Antibody, Alexa Fluor™ 555 | Invitrogen | Cat# A-31572; RRID: AB_162543 |
| Goat anti-Rabbit IgG (H+L) Cross-Adsorbed Secondary Antibody, Alexa Fluor™ 633 | Invitrogen | Cat# A-21070; RRID: AB_2535731 |
| Goat anti-Mouse IgG (H+L) Cross-Adsorbed Secondary Antibody, Alexa Fluor™ 633 | Invitrogen | Cat# A-21050; RRID: AB_2535718 |
| Donkey Anti-Rabbit IgG H&L (Alexa Fluor® 647) preadsorbed | Abcam | Cat# ab150063; RRID: AB_2687541 |
| Goat anti-Mouse IgG (H+L) Cross-Adsorbed Secondary Antibody, Alexa Fluor™ 647 | Invitrogen | Cat# A-21235; RRID: AB_2535804 |
| **Chemicals, peptides, and recombinant proteins** |  |  |
| Vitronectin (VTN-N) Recombinant Human Protein, truncated | Gibco | Cat# A400457 |
| Y-27632 | MCE | Cat# HY-10071 |
| StemPro™-34 SFM (1X) | Gibco | Cat# 10639011 |
| GlutaMAX™ Supplement | Gibco | Cat# 5050061 |
| Human SCF Recombinant Protein | Gibco | Cat# PHC2111 |
| Human Flt-3 Ligand (FLT3L) Recombinant Protein | Gibco | Cat# PHC9414 |
| Human IL-3 Recombinant Protein | Gibco | Cat# PHC0034 |
| Human IL-6 Recombinant Protein | Gibco | Cat# PHC0065 |
| Ethylenediaminetetraacetic acid (EDTA) solution | Millipore Sigma | Cat# E8008 |
| DPBS, no calcium, no magnesium | Gibco | Cat# 14190250 |
| Essential 8™ Medium | Gibco | Cat# A1517001 |
| Penicillin-Streptomycin (10,000 U/mL) | Gibco | Cat# 15140122 |
| POLA1 inhibitor 1 | MCE | Cat# HY-147812 |
| Paraformaldehyde 16% Aqueous Solution EM Grade | EMS | Cat# 15710 |
| Triton™ X-100 | Millipore Sigma | Cat# T8787 |
| PBS, pH 7.4 | Gibco | Cat# 10010023 |
| Fetal Bovine Serum (FBS) | ATCC | Cat# 30-2020 |
| **Hoechst 33342** |  |  |
| Agarose, low gelling temperature | Sigma | Cat# A9045 |
| β-Agarase I | NEB | Cat# M0392 |
| 5-Chloro-2'-deoxyuridine (CldU) | Sigma | Cat# C6891 |
| Idoxuridine (IdU) | MCE | Cat# HY-B0307 |
| MES | Sigma | Cat# M3671 |
| Sodium Chloride | Fisher Scientific | Cat# S271-500 |
| Sodium Hydroxide | Fisher Scientific | Cat# S318-500 |
| TE Buffer (Tris-EDTA Buffer) 1X | Sigma | Cat# 93283 |
| N-lauroylsarcosine sodium salt/Sarkosyl | Sigma | Cat# L9150 |
| Proteinase K, recombinant, PCR grade | Thermo Scientific | Cat# EO0491 |
| Hydrogen peroxide solution | Millipore Sigma | Cat# H1009 |
| RPMI 1640 Medium | Gibco | Cat# 11875093 |
| B-27™ Supplement, minus insulin | Gibco | Cat# 1895601 |
| B-27™ Supplement (50X), serum free | Gibco | Cat# 17504044 |
| Laduviglusib trihydrochloride (CHIR-99021) | MCE | Cat# HY-10182B |
| Wnt-C59 | MCE | Cat# HY-15659 |
| ACCUTASE™ | Stemcell | Cat# 07920 |
| Essential 6™ Medium | Gibco | Cat# A1516401 |
| Alkaline Phosphatase Live Stain | Invitrogen | Cat# A14353 |
| HEPES | Gibco | Cat# 15630080 |
| MgCl<sub>2</sub> | Invitrogen | Cat# AM9530G |
| EGTA, Molecular Biology Grade | Sigma | Cat# 324626 |
| Sucrose solution | Sigma | Cat# 8590 |
| DL-Dithiothreitol solution (DTT) | Sigma | Cat# 646563 |
| NEBuffer™ 2 | NEB | Cat# B7002S |
| Nt.CviPII | NEB | Cat# R0620S |
| DNA Polymerase I (E. coli) | NEB | Cat# M0209S |
| Deoxynucleotide (dNTP) Solution Mix | NEB | Cat# N0447S |
| Aat Bioquest Cyanine 5-dATP [Cy5-dATP] | Aat Bioquest | Cat# 50-224-9676 |
| Protease Inhibitor Cocktail | Sigma | Cat# P1860 |
| TWEEN® 20 | Sigma | Cat# P1379 |
| Nonidet P40 Substitute | Thermo Scientific | Cat# J19628.AP |
| Digitonin (5%) | Invitrogen | Cat# BN2006 |
| RNase Inhibitor | 10X Genomics | Cat# PN-1000887 |
| Buffer RLT Plus | Qiagen | Cat# 1053393 |
| **Critical commercial assays** |  |  |
| CytoTune™-iPS 2.0 Sendai Reprogramming Kit | Invitrogen | Cat# A16517 |
| Click-iT™ Plus EdU Cell Proliferation Kit for Imaging, Alexa Fluor™ 488 dye | Invitrogen | Cat# C10637 |
| Click-iT™ Plus EdU Cell Proliferation Kit for Imaging, Alexa Fluor™ 555 dye | Invitrogen | Cat# C10638 |
| Click-iT™ Plus EdU Cell Proliferation Kit for Imaging, Alexa Fluor™ 594 dye | Invitrogen | Cat# C10639 |
| Click-iT™ Plus EdU Cell Proliferation Kit for Imaging, Alexa Fluor™ 647 dye | Invitrogen | Cat# C10640 |
| Click-iT™ Plus TUNEL Assay Kits for In Situ Apoptosis Detection | Invitrogen | Cat# C10619 |
| Chromium Next GEM Single Cell Multiome ATAC + Gene Expression Reagent Bundle | 10X Genomics | Cat# 1000283 |
| Single Index Kit N Set A | 10X Genomics | Cat# 1000212 |
| Dual Index Kit TT Set A | 10X Genomics | Cat# 1000215 |
| **Deposited data** |  |  |
| GSE351021 |  |  |
| GSE351025 |  |  |
| **Experimental models: Cell lines** |  |  |
| Primary Peripheral Blood Mononuclear Cells (PBMC), Normal, Human | ATCC | Cat# PCS-800-011 |
| hiPSC | Self-derived | N/A |
| Human Cardiomyocyte | Self-derived | N/A |
| **Software and algorithms** |  |  |
| Imaris | Oxford Instruments |  |
| ImageJ | FIJI |  |
| FastQC v0.11.9 |  |  |
| Trimmomatic v0.39 |  |  |
| STAR v2.7.10a | Dobin et al<sup>6</sup> |  |
| SAMtools v1.18 | Li et al<sup>7</sup> |  |
| StringTie v2.2.1 | Pertea et al<sup>8</sup> |  |
| DESeq2 v1.52.0 | Love et al<sup>9</sup> |  |
| deepTools v3.5.4 | Ramirez et al<sup>10</sup> |  |
| IGV v2.16.2 | Robinson et al<sup>11</sup> |  |
| Cellranger-arc count v2.2.0 | Zheng et al<sup>12</sup> |  |
| Seurat v5.5.1 | Hao et al<sup>13</sup> |  |
| Signac v1.17.1 | Stuart et al<sup>14</sup> |  |
| SoupX v1.6.2 | Young et al<sup>15</sup> |  |
| scDBIFinder v1.26.7 | Germain et al<sup>16</sup> |  |
| clusterProfiler v4.20.0 | Xu et al<sup>26</sup> |  |
| CHIPseeker v1.48.0 | Yu et al<sup>27</sup> |  |
| Monocle 3 v1.4.27 | Trapnell et al<sup>28</sup> |  |
| Bioconductor v3.23 | Huber et al<sup>29</sup> |  |
| R v4.6.0 | R Foundation | www.R-project.org |

### **EXPERIMENTAL MODEL**

#### **Cell culture**

The cells were cultured in a humidified 20% O<sub>2</sub>, 5% CO<sub>2</sub> incubator at 37°C. The primary human peripheral blood mononuclear (PBMC) cells were purchased from American Type Culture Collection (ATCC). Cells were maintained in the multi-well cell culture plates (Falcon) in complete PBMC culture medium: StemPro™-34 SFM (Gibco) supplemented with StemPro™-34 Nutrient Supplement (Gibco), 2 mM L-Glutamine (Gibco), 100 ng/ml SCF (Gibco), 100 ng/ml FLT-3 (Gibco), 20 ng/ml IL-3 (Gibco), 20 ng/ml IL-6 (Gibco), 1% Penicillin-Streptomycin (P/S, Gibco). The medium was replaced by half everyday with fresh complete PBMC medium before reprogramming viral transduction. Human induced pluripotent stem cells (hiPSC) were generated and cultured on vitronectin-coated (Gibco) multi-well cell culture-treated plates (Falcon) in Complete Essential 8™ Medium (Gibco). The medium was changed every day. Following passage, the cells were washed in Dulbecco's phosphate-buffered saline (DPBS, Gibco) and incubated for 3 min with Ethylenediaminetetraacetic acid solution (EDTA, Sigma) at 37°C. The cells were collected in a medium with Y-27632 (10 µM, MCE) and pelleted for 5 min at 300 ×g. The cells were resuspended, counted using a TC20 Automated Cell Counter (Biorad), and plated in multi-well plates coated with vitronectin.

### **METHOD DETAILS**

#### **Sendai virus reprogramming**

Human PBMC were reprogramed according to the manufacturer's instructions (CytoTune-iPS 2.0 Sendai Reprogramming Kit, Life technology). Briefly, on day -4, the PBMCs were plated in complete PBMC medium. From day -3 to -1, replace half of the medium with fresh complete PBMC medium. On day 0, cells were transduced with CytoTune™ 2.0 Sendai reprogramming vectors at appropriate MOI. On day 3, transduced cells were plated on vitronectin-coated culture plates in complete StemPro™-34 medium without cytokines: StemPro™-34 SFM medium supplemented with StemPro™-34 Nutrient Supplement, 2mM L-Glutamine, and 1% Penicillin-Streptomycin. From day 4 to 6, replace complete StemPro™-34 medium without cytokines every day. On day 7, Essential 8™ Medium was used to replace half of the complete StemPro™-34 medium without cytokines to start transition into full Essential 8™ Medium to generate passage 1 (P1) hiPSC. From Day 8 up to 3 weeks, medium was replaced fully with Essential 8™ Medium every day to continue culturing cells on vitronectin-coated culture dishes. hiPSC colonies were manually picked, passaged, and cultured based on morphology and colony density.

#### **POLA1 inhibitor 1 treatment**

POLA1i inhibitor 1 (PolA1i, MCE) was purchased and diluted into working concentration by medium used for the cell culture. For reprogramming, PolA1i was added to the reprogramming medium with a final concentration of 10 nM from Day -2 to Day 3 replenished daily.

#### **Immunofluorescence Imaging**

For day 0 and day 3 samples, PBMCs were centrifuged onto precoated plates at $450 \times g$ for 5 min. Before use, the plates were coated with 0.1 mg/mL poly-D-lysine hydrobromide (Sigma-Aldrich, P7280) for 90 min at room temperature and washed twice with PBS (Gibco). Following attachment to the plates, the cells were processed for immunofluorescence staining as described above. Cells were fixed with 4% paraformaldehyde (EMS) for 20 minutes at room temperature. Subsequently, an optional permeabilization step was performed only for intracellular marker detection using PBS supplemented with 0.2% (v/v) Triton X-100 (Sigma-Aldrich). Then, cells were washed with PBS before blocking using PBS supplemented with 0.2% (v/v) Triton X-100 (0.5% for intracellular marker detection) and 5% (w/v) bovine serum albumin (BSA, ATCC) for 1 hour at room temperature with gentle shaking. Primary antibodies were diluted to appropriate concentrations using blocking solution, and incubated with the cells overnight at 4°C. Following primary antibody incubation, the cells were washed with PBS for 15 minutes for three times. Then, the cells were incubated with secondary antibodies in blocking solution for 2 hours at room temperature. Nuclear staining was completed using Hoechst 33342 (1:1000). Images were acquired on a Leica STELLARIS confocal laser scanning microscope equipped with LIGHTNING detection using a $\times 63$ Apo NA1.3 oil immersion objective (Leica).

#### **DNA Fibers in PBMC**

DNA fibers were prepared as described<sup>1</sup>. PBMC samples during reprogramming time points were subsequently pulse labeled with 25 μM 5-iodo-2'-deoxyuridine (IdU; MCE) and 250 μM 5-chloro-2'-deoxyuridine (CldU; Sigma-Aldrich) for 1 hour each and collected. The labeled cells were embedded into the 5% Low-melting-point (LMP) agarose to be stored as plugs. The cells in the plugs were lysed in lysing buffer supplemented with Proteinase K (Fisher) in 50°C water bath overnight. On day 2, the plugs were washed with TE buffer and treated with β-Agarase at 42°C overnight to release DNA fibers from the plug into the solution. By using the Fiber Comb machine (Genomic Vision), the silanized glass coverslip (PolyAn) was submerged into the DNA fibers solution and removed at a slow constant rate (300 mm/s), and each DNA fiber was straightened and aligned along the same axis. The DNA fibers were then dried in 65°C incubator for 2 hours and denatured in denature solution consisting of 0.5 M NaOH (Fisher) and 1 M NaCl (Fisher). Stepwise de-hydration was performed from 70% ethanol to 100% ethanol followed by air-dry over 30 minutes. The subsequent blocking was completed using PBS supplemented with 5% (w/v) BSA in 37°C incubator for 1 hour. CldU and IdU fiber signals were detected with anti-bromodeoxyuridine (anti-BrdU) antibodies recognizing CldU (BD<sup>TM</sup>) and IdU (Abcam), respectively, and appropriate secondary antibodies. After the detection of IdU and CldU tracks, DNA was detected using antibody against single-stranded DNA (ssDNA, Sigma). Images were acquired as describe above using Leica STELLARIS confocal laser scanning microscope. To calculate fork speed, we used the established conversion 1 μm = 2 kb as described<sup>2</sup>. Analysis of DNA fibers was performed by using a manual processing pipeline that consisted of three steps: (1) categorization of replication pattern, (2) quantification of fiber lengths via Fiji (ImageJ), and (3) statistical analysis of different fiber parameters (such as replication pattern and speed). Briefly, for all images acquired, the same Macro was run to process the images to the same intensity range and global scale was set for the images for later quantification. Line drawing tool was utilized within Fiji facilitated by an external drawing pad to accurately measure the length of each track (green/red) of each fiber. Only bi-colored fibers overlapping with DNA counterstaining were included. There was a minimum of 100 fibers measured for each sample.

#### **Histone Co-localization Assay**

Cells were pulse-labeled with EdU for 25 min. Following sample collection, soluble nuclear proteins were removed by incubating the cells for 2 min at room temperature with ice-cold clearance buffer containing 10 mM HEPES (Gibco), 100 mM NaCl, 3 mM MgCl<sub>2</sub> (Invitrogen), 1 mM EGTA (Sigma), 300 mM sucrose (Sigma), 0.2% (v/v) Triton X-100, and 0.2% (w/v) BSA. Immediately before use, the buffer was supplemented with 1x protease inhibitor cocktail (Sigma) and 1 mM DTT (Sigma). Following clearance, cells were fixed with 4% PFA and processed for immunofluorescence staining as described above. Images were acquired as describe above using Leica STELLARIS confocal laser scanning microscope in LIGHTENING mode. The images were processed in Imaris software (Oxford Instruments) to exclusively crop the EdU-positive cells at single cell level that are at early to mid-S phase stage, then the Coloc function in Imaris was used with masking of DNA or EdU signal to find the Pearson's Correlation Coefficient between corresponding histone modification signal pairs. The data was recorded for statistical analysis. Sample group size is $n > 29$.

#### **EdU incorporation assay**

A 2X working solution of EdU (Thermo) was prepared following the manufacture's instruction of Click-iT™ Plus EdU Cell Proliferation Kit for Imaging (Invitrogen). The solution was prepared in complete PBMC medium from the 10mM stock with a final treatment concentration of 10 μM. The cells were pulse labeled by EdU for 1 hour. The final samples were process following immunofluorescence process as described above. Finally, Click-iT® reaction was performed following the manufacture's instruction (Invitrogen) for EdU signal detection. Images were acquired as describe above using Leica STELLARIS confocal laser scanning microscope. The images were processed and quantified in ImageJ software.

#### **NicE-View Assay**

Cells were processed as described for immunofluorescence. Following primary antibody incubation, cells were washed three times with 0.2% PBST (0.2% (v/v) TWEEN® 20 in PBS), and washed once with 1× NEBuffer (NEB) and incubated in the buffer for 5 min at room temperature. Cells were then incubated with accessible chromatin labeling buffer containing 1× NEBuffer, 2.5 U Nt.CviPII nicking endonuclease (NEB), 50 U DNA

Polymerase I (NEB), and 30 μM total dNTPs (NEB), including 6 μM Cyanine 5-dATP (Cy5-dATP; AAT Bioquest). Labeling was performed at 37°C for 90 minutes. The reaction was quenched by replacing the labeling buffer with 50 mM EDTA in 0.2% PBST. After incubation for 5 minutes at room temperature, cells were washed twice with 0.2% PBST and imaged following method as described above.

#### **TUNEL assay**

TUNEL assay was performed with Click-iT™ Plus TUNEL Assay for In Situ Apoptosis Detection (Invitrogen) according to the manufacturer's instructions (Invitrogen). Briefly, the samples were incubated with TdT Reaction Buffer for 10 minutes at 37°C, and then incubated with TdT reaction mixture for 60 minutes at 37°C. After washing with 5% (w/v) BSA and 0.5% (v/v) Triton X-100 in PBS for three times, the samples were incubated with the Click-iT® Plus TUNEL reaction cocktail at 37°C for 30 minutes, protected from light. Finally, the samples were washed with 5% (w/v) BSA in PBS for three times followed by immunostaining and imaging as described above.

#### **UVA irradiation stress assay**

An UVA light source (wavelength: 315-400 nm) was set up in a 36°C incubator and calibrated by a UV dosimeter with irradiance intensity of 8 mW/cm<sup>2</sup>. The live cell samples were washed with DPBS three times and kept in DPBS before and during UVA irradiation treatment. The cell plates lid was removed for full exposure of UVA light source and positioned at a 10 cm distance from the light source and irradiated for 3 minutes 15 seconds to deliver a total dose of 3 J/cm<sup>2</sup>. The control cells were kept in DPBS and incubated in the same incubator condition in dark during the irradiation period. The samples were washed by DPBS followed by replenishment of complete media immediately after UVA irradiation.

#### **H<sub>2</sub>O<sub>2</sub> oxidative stress assay**

Cells were treated with 20-50 μM H<sub>2</sub>O<sub>2</sub> (Sigma) for 1 hour under standard culture conditions. Following treatment, cells were washed with DPBS three times and maintained in fresh culture medium for subsequent analysis.

#### **Cardiomyocyte differentiation from hiPSCs**

The hiPSCs were differentiated into cardiomyocytes as described previously<sup>3</sup>. Briefly, on day 0, the cells were incubated for 1 day with RPMI-1640 (Gibco) supplemented with 2% B27 supplement without insulin (Gibco), 6 μM CHIR99021 (MCE), and 1% P/S. On day 1, the cells were rinsed with DPBS and incubated in RPMI-1640 supplemented with B27 without insulin. On day 2 and day 3, the cells were incubated with RPMI-1640 supplemented with B27 without insulin and 5 μM C59 (MCE). From day 4 to day 6, the cells were washed with DPBS and incubated with RPMI-1640 supplemented with B27 without insulin. Then from day 6 to day 8, the cells were incubated with RPMI-1640 supplemented with B27 with insulin. From day 8, the differentiated cardiomyocytes are maintained in RPMI-1640 supplemented with B27 with insulin.

#### **Embryoid body formation and Spontaneous differentiation from hiPSCs**

On day 0, iPSCs were dissociated by Accutase (Stemcell) into small aggregates and transferred to low-attachment V-bottom 96-well plates (Corning) to promote embryoid body formation. EBs were maintained in suspension culture in complete hiPSC Essential 8™ Medium replenished daily from day 0 to day 4. On day 4, half of complete hiPSC medium was replaced by Essential 6™ Medium (E6, Gibco). From day 5, half volume media change was completed by E6 Medium every day until day 12. Daily bright field images were recorded. On day 12, embryoid bodies were processed following immunofluorescence protocol as described above with prolonged permeabilization (1 hour), blocking (overnight), and primary antibody incubation period (24 hours).

#### **RNA-seq of reprogrammed iPSC colonies**

Emerging iPSC colonies during PBMC reprogramming were cultured in Essential 8 medium (Gibco) until day 13, 17, or 21. At each time point, colonies from the different treatment conditions were manually picked and lysed directly in 10 µL Buffer RLT Plus (Qiagen). Poly(A) RNA was captured with biotinylated oligo-dT primers, and libraries were prepared following the G&T-seq protocol<sup>4</sup>. Libraries that passed quality control were pooled and sequenced (paired-end, 150 bp) on a NovaSeq X Plus instrument (Novogene Corporation, CA, USA).

Raw FASTQ files were assessed with FastQC (v0.11.9), and adapters were trimmed with Trimmomatic (v0.39) using the Nextera adapter sequences<sup>5</sup>. Trimmed read pairs were aligned to the human reference genome (GRCh38, Ensembl release 114, with the matching GTF annotation) using STAR (v2.7.10a)<sup>6</sup>. Alignments were indexed with SAMtools (v1.18)<sup>7</sup>. Transcript assembly and quantification, including generation of the gene count matrix, were performed with StringTie (v2.2.1)<sup>8</sup>. Counts were normalized with DESeq2 (v1.52.0)<sup>9</sup>. Normalized expression of genes of interest was visualized as heatmaps. Pairwise Euclidean distances between samples were calculated from the normalized data to assess sample similarity. To visualize SUSD2 and H3Y1 read coverage, BigWig files were generated with deepTools (v3.5.4)<sup>10</sup> and averaged across samples within each condition, then displayed in IGV (v2.16.2)<sup>11</sup>.

#### **Single cell multi-omic sample preparation**

For day 0 (D0) and day 3 (D3) samples, PBMCs were collected and centrifuged at 500 × g for 5 min at room temperature. For day 7 (D7) samples, adherent cells were dissociated using Accutase (Stemcell), followed by the addition of an equal volume of StemPro medium (Gibco) to quench dissociation. Cells were then pelleted by centrifugation at 500 × g for 5 min at room temperature. All samples were washed once with ice-cold PBS containing 0.04% BSA, followed by centrifugation at 500 × g for 5 min at 4°C. The supernatant was removed, and cell pellets were resuspended in 50 µL of ice-cold lysis buffer containing 10 mM Tris-HCl (pH 7.4), 10 mM NaCl, 3 mM MgCl<sub>2</sub>, 0.1% Tween-20 (Sigma), 0.1% Nonidet P40 Substitute (Thermo), 0.01% digitonin (Invitrogen), 1% BSA (ATCC), and 1× RNase inhibitor (10X Genomic) in nuclease-free water. D0 and D3 samples were lysed on ice for 3 min, whereas D7 samples were lysed on ice for 4 min. Lysis was quenched by adding 50 µL of wash buffer containing 10 mM Tris-HCl (pH 7.4), 10 mM NaCl, 3 mM MgCl<sub>2</sub>, 0.1% Tween-20, 1% BSA, and 1× RNase inhibitor in nuclease-free water. Nuclei were pelleted by centrifugation at 500 × g for 5 min at 4°C, and the supernatant was removed. Subsequently, 50 µL of chilled diluted

Nuclei Buffer was gently added to the pellet without mixing, followed by centrifugation at $500 \times g$ for 5 min at 4°C. The supernatant was removed, and nuclei were resuspended in 7 $\mu\text{L}$ of diluted Nuclei Buffer, of which 2 $\mu\text{L}$ was used for counting. A total of ~10,000 nuclei per sample were loaded onto the Chromium platform, and libraries were prepared using the Chromium Next GEM Single Cell Multiome ATAC + Gene Expression Reagent Bundle (10X Genomics) according to the manufacturer's instructions. ATAC libraries were indexed using the Single Index Kit N Set A (10X Genomics), and gene expression libraries were indexed using the Dual Index Kit TT Set A 96 (10X Genomics).

#### Single cell multi-omic data analysis

The demultiplexed FASTQ files were processed with “**cellranger-arc count**” (10x Genomics, v2.2.0)<sup>12</sup> using the human reference (GRCh38) - 2024-A downloaded from 10xGenomics whose outputs were analyzed using standard pipelines of the **Seurat** (v5.5.1)<sup>13</sup> and **Signac** (v1.17.1)<sup>14</sup> R packages. The single cell RNAseq data for each sample were first checked with **SoupX** (v1.6.2)<sup>15</sup> to remove ambient RNA contamination where 0.15 is set as the sanity ceiling. After that, single nuclei passed these RNAseq criteria: $nFeature\_RNA > 200$ & $nFeature\_RNA < 15000$ & $nCount\_RNA > 1000$ & $nCount\_RNA < 50000$ & $percent.mt < 20$ were used to construct a Seurat object and **scDblFinder** (v1.26.7)<sup>16</sup> was used to distinguish and remove doublets. Finally, these criteria: $nCount\_ATAC < 100000$ & $nCount\_RNA < 30000$ & $nCount\_ATAC > 1800$ & $nCount\_RNA > 1000$ & $nucleosome\_signal < 2$ & $TSS.enrichment > 1$ were used to further subset the singlets. All nuclei in each sample that passed all above filtering procedures were used to create the scRNAseq Seurat object and scATACseq Seurat object. Both Seurat objects were normalized and stored for downstream analyses.

Saved Seurat objects from scRNAseq and scATACseq of each sample were merged for joint analysis. UMAP was generated by Seurat with resolution set to 0.8. The markers of PBMC lineages<sup>17–23</sup> and iPS cells<sup>24,25</sup>, as well as the cell annotation by cellranger were applied to distinguish different cell types. The FindMarkers function of Seurat was used for differential analysis using the pseudo-bulked RNAseq data. Transcripts that 20% changes in the PolA1i treated samples than the corresponding control samples and had adjusted p values $< 0.1$ in the Benjamini-Hochberg (BH) test were considered differentially expressed which were further used for gene ontology (GO) and gene set enrichment analysis (GSEA). GO and GSEA was performed and visualized with the **clusterProfiler** (v4.20.0)<sup>26</sup> package. **ChIPseeker** (v1.48.0)<sup>27</sup> was used to annotate the ATACseq peaks captured in each sample. The scRNAseq and scATACseq data were used jointly to find and label different clusters, and the results were visualized by UMAP, after which **monocle 3** (v1.4.27)<sup>28</sup> were used for pseudotime analysis. The CellCycleScoring function of Seurat was used to explore the cell cycle phases of nuclei. **Bioconductor** (v3.23)<sup>29</sup> and **R** (v4.6.0, www.R-project.org) as well as other necessary packages were used to finish all analysis tasks.

#### STATISTICAL ANALYSIS

All experiments used PBMCs from 6 independent donors. For all analyses, a biological replicate was defined as an independent donor. Wells, imaging fields, nuclei, DNA

fibers, and individual colonies within a replicate were treated as technical units and were not counted as independent samples, except where explicitly stated. The numbers of donors, independent experiments, wells, colonies, and iPSC clones analyzed for each panel are given in the figure legends. Data are presented as mean ± SEM unless otherwise indicated. Comparisons between two groups used two-tailed unpaired / paired Student's t-test or Mann–Whitney U test, and comparisons among more than two groups used one-way ANOVA with Tukey's / Dunnett's post hoc test. Experiments with two factors (treatment and time point) were analyzed by two-way ANOVA with Šidák's correction. Fold changes relative to control were analyzed on the log scale using a one-sample t-test or a paired test. Proportions (e.g., established versus failed iPSC lines) were compared using Fisher's exact test. For imaging measurements with many nuclei per replicate, data were analyzed using nested or mixed-effects models with donor/experiment as a random effect, or after averaging within each replicate. For snRNA-seq and snATAC-seq data, differential comparisons used a Wilcoxon rank-sum test with Benjamini–Hochberg correction. P values are indicated as \*P < 0.05, \*\*P < 0.01, \*\*\*P < 0.001, \*\*\*\*P < 0.0001, and ns, not significant. Colony counts and imaging quantification were performed by automated analysis. Statistical analysis was performed with GraphPad Prism v10.6.1(892).
